# Measuring changes in transmission of neglected tropical diseases, malaria, and enteric pathogens from quantitative antibody levels

**DOI:** 10.1101/106708

**Authors:** Benjamin F. Arnold, Mark J. van der Laan, Alan E. Hubbard, Cathy Steel, Joseph Kubofcik, Katy L. Hamlin, Delynn M. Moss, Thomas B. Nutman, Jeffrey W. Priest, Patrick J. Lammie

**Affiliations:** School of Public Health, University of California, Berkeley, CA, USA; Laboratory of Parasitic Diseases, National Institute of Allergy and Infectious Diseases, National Institutes of Health, Bethesda, MD, USA; Division of Parasitic Diseases and Malaria, United States Centers for Disease Control and Prevention, Atlanta, GA, USA; Division of Foodborne, Waterborne, and Environmental Diseases, United States Centers for Disease Control and Prevention, Atlanta, GA, USA; Neglected Tropical Diseases Support Center, Task Force for Global Health, Decatur, GA, USA

## Abstract

**Background:** Serologicalantibody levels are a sensitive marker of pathogen exposure, and advances in multiplex assays have created enormous potential for large-scale, integrated infectious disease surveillance. Most methods to analyze antibody measurements reduce quantitative antibody levels to seropositive and seronegative groups, but this can be difficult for many pathogens and may provide lower resolution information than quantitative levels in low transmission settings. Analysis methods have predominantly maintained a single disease focus, yet integrated surveillance platforms would benefit from methodologies that work across diverse pathogens included in multiplex assays.

**Methods/Principal Findings:** We developed an approach to measure changes in transmission from quantitative antibody levels that can be applied to diverse pathogens of global importance. We compared age-dependent immunoglobulin G curves in repeated cross-sectional surveys between populations with differences in transmission for multiple pathogens, including: lymphatic filariasis (*Wuchereria bancrofti*) measured before and after mass drug administration on Mauke, Cook Islands, malaria (*Plasmodium falciparum*) before and after a combined insecticide and mass drug administration intervention in the Garki project, Nigeria, and enteric protozoans (*Cryptosporidium parvum*, *Giardia intestinalis*, *Entamoeba histolytica*), bacteria (enterotoxigenic *Escherichia coli*, *Salmonella spp*.), and viruses (norovirus groups I and II) in children living in Haiti and the USA. Age-dependent antibody curves fit with ensemble machine learning followed a characteristic shape across pathogens that aligned with predictions from basic mechanisms of humoral immunity. Differences in pathogen transmission led to shifts in fitted antibody curves that were remarkably consistent across pathogens, assays, and populations. Mean antibody levels correlated strongly with traditional measures of transmission intensity, such as the entomological inoculation rate for *P. falciparum* (Spearman’s rho=0.75). Seroprevalence estimates recapitulated patterns observed in quantitative antibody levels, albeit with lower resolution.

**Conclusions/Significance:** Age-dependent antibody curves and summary means provided a robust and sensitive measure of changes in transmission, with greatest sensitivity among young children. The method generalizes to pathogens that can be measured in high-throughput, multiplex serological assays, and scales to surveillance activities that require high spatiotemporal resolution. The approach represents a new opportunity to conduct integrated serological surveillance for neglected tropical diseases, malaria, and other infectious diseases with well-defined antigen targets.

**Author Summary:** Global elimination strategies for infectious diseases like neglected tropical diseases and malaria rely on accurate estimates of pathogen transmission to target and evaluate control programs. Circulating antibody levels can be a sensitive measure of recent pathogen exposure, but no broadly applicable method exists to measure changes in transmission directly from quantitative antibody levels. We developed a novel method that applies recent advances in machine learning and data science to flexibly fit age-dependent antibody curves. Shifts in age-dependent antibody curves provided remarkably consistent, sensitive measures of transmission changes when evaluated across many globally important pathogens (filarial worms, malaria, enteric infections). The method’s generality and performance in diverse applications demonstrate its broad potential for integrated serological surveillance of infectious diseases.

## Introduction

There is large overlap in the distribution of global disease burdens attributable to neglected tropical diseases (NTDs), malaria, enteric infections and under-vaccination. Despite nearly a decade of advocacy for integrated monitoring and control [1], prevailing surveillance efforts maintain a single-disease focus, and the high cost of fielding surveys to collect specimens means that programs conduct surveillance infrequently or not at all. High throughput, multiplex antibody assays enable the simultaneous measurement of quantitative antibody responses to dozens of pathogens from a single blood spot [2]. When coupled with existing surveillance platforms, multiplex antibody assays could enable the global community to more quickly identify public health gaps, including: recrudescence of NTD or malaria transmission in elimination settings, stubborn areas of high transmission, emerging infectious diseases, and under-vaccination. Of particular interest are methods to analyze measurements collected in cross-sectional surveys because most large-scale global surveillance efforts use this design (e.g., immunization coverage surveys, malaria indicator surveys, transmission assessment surveys for NTD elimination programs, demographic and health surveys).

A unique attribute of antibody measurements is that they provide an immunological record of an individual’s exposure or vaccination history, and thus integrate information over time [3]. Yet, the information contained in circulating antibodies varies greatly by pathogen and antibody measured, and it is this complexity that presents challenges to the use of antibody measurements for integrated surveillance. Most previous studies have reduced quantitative antibody measurements to seropositive and seronegative groups by choosing a cut point, and then have used models to estimate seroconversion rates from age-dependent seroprevalence as a measure of pathogen transmission [3,4]. The choice of seropositivity cut point can be ambiguous for many pathogens, as examples in this article will illustrate, and can vary widely in lower transmission settings depending on the reference population or statistical method used [5]. A second challenge in lower transmission settings is that seropositive individuals are extremely rare, and so accurate estimates of seroprevalence require large samples [6]. If quantitative antibody levels provide similar information to seroprevalence, relying on quantitative levels directly could enable surveillance efforts to detect differences in populations with smaller samples or at higher resolution. Thus, analytical methods that use the quantitative response directly avoid the difficulty of defining cut points, naturally accommodate gradual waning of immunity with age that can present difficulties to seroconversion models [4], and may provide higher resolution information in lower transmission settings.

To our knowledge there has not been a broad-based assessment for whether quantitative antibody measurements present an opportunity for integrated surveillance across diverse pathogens. Two recent contributions in the malaria literature proposed mathematical models to measure changes in transmission from quantitative antibody responses [7,8]. Both models require strong parametric assumptions such as constant rates of antibody acquisition and loss over different ages, or constant transmission over time, which may be difficult to justify for many pathogens of interest in an integrated surveillance platform.

Our objective was to develop a general and parsimonious method to measure changes in infectious disease transmission from quantitative antibodies. We approached the problem from a different perspective than mathematical modeling, and instead focused on recent advances in machine learning and statistical estimation theory to measure differences in transmission within or between populations. We also aimed to assess whether the method could generalize across diverse pathogens that can be measured in multiplex assays, such as neglected tropical diseases, malaria, and enteric pathogens. A widely observed phenomenon across infectious diseases is that changes in pathogen transmission result in a “peak shift” of infection intensity by age: as transmission intensity declines in a population, the age-specific prevalence and intensity of infection tends to rise more slowly at younger ages and peak at lower overall levels [9]. We sought to extend this observation to measure changes in transmission using quantitative antibody levels rather than measures of patent infection -- an approach suggested by mathematical models of parasite immunity [9,10] with empirical support in a comparison of populations with varying helminth transmission intensity [11].

We focused on a general mechanism of acquired immunity elicited by most infectious pathogens. Children are born with maternal immunoglobulin G (IgG) antibodies that wane over the first 3-6 months of life, and from ages 4-6 weeks begin to produce their own IgG antibodies in response to antigen exposure [12]. The aggregation of individual IgG responses generates a curve of population average IgG levels that rises in the first years of life until it plateaus at adult levels [13]. Transferred maternal immunity -- a function of maternal immunologic memory -- likely influences the magnitude of the population-average IgG curve’s intercept near birth [12]. Antigen exposure is needed to maintain antibodies in blood, either by stimulating the proliferation of memory B-cells to replenish short-lived plasma cells or by stimulating the production of non-germinal center short-lived plasma cells [13]. Antigen exposure induces rapid proliferation and differentiation of short-lived B-cells, with somatic hypermutation leading to increased affinity following each exposure. As transmission declines, population-average serum IgG levels should rise more slowly as the age of first infection increases and repeated exposures become infrequent. For pathogens that elicit antibody responses that wane over time, the number of long-lived antibody secreting cells should decline without recent antigen exposure [13], which in turn should be reflected in a lower plateau of the age-dependent antibody curve. We therefore hypothesized that reduced pathogen transmission would cause pathogen-specific IgG antibody curves to increase more slowly with age and plateau at lower levels, and that quantifying changes in the curves would provide a robust and sensitive measure of changes in transmission within or between populations.

## Methods

### Overview of the approach

To test this hypothesis, we examined age-dependent antibody responses (“age-antibody curves”) to diverse pathogens in populations with likely differences in transmission intensity. We fit age-antibody curves with a data adaptive, ensemble machine learning algorithm that can include additional covariates to control for potential confounding [14]. The curves represent a predicted mean antibody level by age (*a*) for each exposure group (*x*), which we denote *E*(*Y*_*a,x*_) in the statistical methods. We used the age-adjusted mean antibody response within each group (*x*) as a summary measure of transmission, denoted *E*(*Y*_*x*_), and estimated differences between group means. The age-adjusted mean antibody response equals the area under the age-antibody curve (Text S1). The approach thus integrates the steepness of the curve’s initial rise at young ages as well as its sustained magnitude at older ages, with lower transmission measured by reductions in group means. Comparing group means intuitively represents an average difference between groups across all points in the curves. If particular age ranges are of interest, such as young children, then the mean can be estimated over restricted regions of the age-antibody curve.

### Lymphatic filariasis transmission on Mauke Island

Mauke, Cook Islands was endemic for *Wuchereria bancrofti* in decades past, and in 1987 there was an island-wide MDA of all individuals ≥5 years old with diethylcarbamazine. The present analysis included serum samples from two cross-sectional measurements of the permanent resident population; the first in 1975 (N=362, approximately 58% of the population) and the second in 1992, 5 years after the island-wide MDA (N=553, approximately 88% percent of the population) [15]. Both studies preserved serum samples by freezing them in liquid nitrogen within hours of collection and storing them at -80℃. Serum samples were tested for IgG antibody levels to the Wb123 antigen using a Luciferase Immunoprecipitation System (LIPS) assay, as previously described in detail [16]. Data presented are in luminometer units from averaged duplicate samples.

We re-analyzed data from the original assessment of the effect of the MDA campaign on Wb123 antibody levels [15] using the statistical methods described below. We estimated separate age-antibody curves in 1975 and 1992. To make statistical comparisons between the curves, we estimated means for each survey year and differences between surveys, stratified by 5 year age group for ages ≤20 years old. For a subsample of 114 individuals who were measured in both 1975 and 1992, we compared Wb123 antibody levels in subgroups defined by whether they had circulating antigen to adult *W. bancrofti* -- an indication of active infection -- at one or both time points. We plotted individual changes in Wb123 antibody levels to visualize antibody acquisition and loss in different subgroups.

### Malaria transmission in the Garki Project, Nigeria

The Garki Project, led by the World Health Organization and the Government of Nigeria, included a comprehensive malaria intervention study that took place in 22 villages in the rural Garki District, Nigeria (1970-1976) [17]. We obtained publicly available study datasets for this analysis (http://garkiproject.nd.edu). The intervention included a combination of insecticide spraying and mass drug administration of surfanene-pyrimethamine in 1972-1973, along with targeted distribution of chloroquine to children <10 and self-reporting fever cases in the 1974-75 post-intervention period. The study documented large reductions in the proportion of individuals testing positive for *Plasmodium falciparum* infection by microscopy as a result of the intervention.

In a subset of two control villages and six intervention villages, the study collected multiple serological measures that have been described in detail [17]. Briefly, the study collected serum from all members present in a village in eight rounds that alternated between wet and dry seasons. We limited the analysis to 4,774 specimens collected from individuals <20 years old because that age range captured nearly all of the change in the age-antibody curve (median serum samples per round in each village: 74, range: 19-158). Serological survey rounds 1-2 took place in the wet and dry season before the intervention started, rounds 3-5 took place during the active intervention period at 20, 50, and 70 weeks after intervention initiation, and rounds 6-8 took place at 20, 40, and 90 weeks after the conclusion of intervention activities. The sixth measurement was collected in the intervention villages only. From each participant, finger prick blood samples were collected in two 0.4-ml heparinized Caraway tubes for immunological testing. We focused on *P. falciparum* antibody response measured with the IgG indirect fluorescent antibody (IFA) test. We converted IFA titers to the log_10_ scale and then estimated mean IFA titre by age separately for intervention and control villages in each measurement round. We compared curves using the difference between age-adjusted means. We repeated the analysis at the village level to make separate comparisons of each individual intervention village against control to examine curves and measures of transmission at smaller spatial scale.

The study collected extensive wet season entomological measurements in three of the villages with serological monitoring. The co-located entomological and serological measurements enabled us to compare village-level mean antibody levels and seroprevalence with the wet season entomological inoculation rate (EIR) as the transmission intensity changed in the intervention villages. EIR estimates from Table 4 of the original study [17] were used in the analysis. The EIR represents the number of sporozoite positive bites per person over each wet season, and was estimated by multiplying the man-biting rate by the sporozoite positive rate in night-bite collections. Night-bite collections were conducted every 2 weeks using 2 indoor and 2 outdoor stations per village, with 2 human bait collectors in each station throughout the night. We estimated village level mean IFA antibody titers restricted to serum samples collected during the same periods of EIR monitoring, and we measured the association between village level mean antibody titers and the EIR with the Spearman rank correlation coefficient.

After completing the primary analysis that estimated age-antibody curves by survey for control and intervention villages, we noticed a reduction in age-adjusted geometric mean antibody titers between wet and dry survey rounds 1-2. We followed-up this observation with a secondary analysis, restricted to the control villages, that estimated age-antibody curves separately by survey round, which corresponded to wet and dry seasons: 1971 wet (survey 1), 1972 dry (survey 2), 1972 wet (survey 3), 1973 dry (survey 4), and 1973 wet (survey 5) [17]. Control villages were not measured in survey 6, and surveys 7-8 took place in the 1974 and 1975 wet seasons; we excluded surveys 7-8 from the secondary analysis because we were interested in comparing transmission in adjacent wet and dry seasons.

## Enteric pathogen transmission in Haiti and the United States

Our analysis of enteric pathogen antibody measurements relied on two existing data sources. Haiti samples were collected from a longitudinal cohort of 142 children, enrolled between the ages of 1 month and 6 years on a rolling basis from 1991-1999 to monitor lymphatic filariasis, and the selection of samples from the Haiti cohort has been described in detail [18]. Children were followed up to 9 years (median 5 years) and each child was measured approximately once per year. At each measurement, the study collected finger prick blood samples. The multiplex bead assay techniques and antibody results for the *Cryptosporidium parvum* recombinant 17-kDa and 27-kDa antigens [19], the VSP-5 fragment of *Giardia intestinalis* variant-specific surface protein 42e [20], and the *Entamoeba histolytica* lectin adhesion molecule (LecA) [21] have been described [18,22]. Enterotoxigenic *Escherichia coli* (ETEC) heat labile toxin *β* subunit [23] and lipopolysaccharide (LPS) from *Salmonella enterica* serotype Typhimurium (Group B) [24] were purchased from Sigma-Aldrich (St. Louis, MO). Purified recombinant norovirus GI.4 and GII.4 New Orleans [25] virus-like particles from a baculovirus expression system [26] were kindly provided by J. Vinje and V. Costantini (CDC, Atlanta, GA). Proteins and LPS were coupled to SeroMap beads (Luminex Corp. Austin, TX) at 120 μg per 12.5 x 106 beads in phosphate-buffered saline at pH 7.2 and were included in the multiplex bead assays previously described [18].

As part of a serologic study in the United States (USA) [27], our lab (JWP, PJL) had banked 86 anonymous blood lead samples collected in 1999 from children ages 0-6 years. The USA samples were tested contemporaneously with the Haiti longitudinal cohort using the same techniques and bead preparations [18]. We used these anonymous samples from the USA to compare antibody curves with the Haitian children.

For each enteric antibody, we estimated separate age-antibody curves in the USA and Haiti using all measurements collected at ages <5.5 years (ages of overlap between the sample sets). We then estimated geometric means for each population and differences between means as described in the statistical methods.

### Statistical methods

A cross-sectional survey measures an individual’s quantitative antibody level (*Y*), age (*A*), and other characteristics (*W*). Many surveillance efforts are also interested in differences in antibody levels by one or more exposures (*X*), which could be confounded by *A* and *W*. We assumed the observed data *O=* (*Y, A, W, X*) ~ *P*_*0*_ arose from a simple causal model (Text S2 includes additional details): *W* = *f*_*W*_ (*U*_*W*_); *A* = *f*_*A*_ (*U*_*A*_); *X* = *f*_*X*_ (*A, W, U*_*X*_); *Y* = *f*_*Y*_(*X, A, W, U*_*Y*_).

### Estimation of age-dependent antibody curves

We estimated the mean antibody level by age, conditional on exposure to *X, E*(*Y*_*a,x*_), and potentially adjusted for covariates *W*:

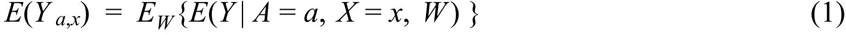

The functional form between antibody response and age can be very nonlinear and differ depending on the population, pathogen, and antibody studied. Data adaptive machine learning provides a robust and flexible estimation approach for the curves [4]. We used an ensemble algorithm called “super learner” that uses cross-validation to combine many different algorithms into a single prediction [14]; the ensemble prediction has cross-validated prediction error less than or equal to any of its constituent algorithms. Ensemble approaches are particularly useful for applications like integrated surveillance when no single model or algorithm will consistently provide the best fit to the data across pathogens and populations. Including a diverse library of models and algorithms in the ensemble ensures the best estimation of the age-antibody relationship across diverse applications. We fit age-dependent antibody curves using the ensemble, and then estimated the marginally adjusted antibody level for each age in the observed data (additional details in Text S2).

### Summary of the curve

We targeted the mean antibody response adjusted for age (*A*) and potential covariates (*W*), conditional on exposure group (*X*=*x*):

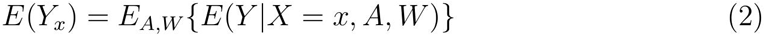

The mean antibody response is a smooth functional of the curve, which makes it tractable and efficient to estimate from a statistical perspective. The mean equals the area under the age-antibody curve (AUC) when age is independent of covariates *W*; if the independence assumption does not hold, then it is equivalent to estimating the AUC within strata of *W* (Text S1). The mean (equal to AUC) is a useful summary measure because it incorporates both the steepness of the initial rise in the curve at early ages as well as the sustained height of the curve at older ages -- for these same reasons this summary measure is widely used for vaccine response in individuals over time and in bioequivalence studies [28].

We compared mean antibody levels in populations with different levels of exposure (*X*), for example comparing Wb123 antibody levels to *W. bancrofti* before (*X*=0) versus after (*X*=1) mass drug administration. Our target parameter of interest was the marginal difference between groups in mean antibody levels, averaged over age (*A*), and potentially confounding covariates (*W*):

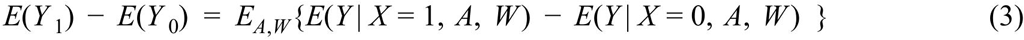

We estimated the difference in mean antibody levels using with targeted maximum likelihood estimation (TMLE) [29]. TMLE is a double-robust, efficient estimation approach that naturally incorporates machine learning in the estimation process and still recovers accurate statistical inference [30]. We estimated influence curve-based standard errors and *P*-values, which accounted for repeated observations if that was a feature of the design [29]. Stratified analyses (e.g., village estimates or estimates within age bands) stratified the data, estimated the mean difference within strata, and then adjusted *P*-values using a Bonferroni correction (additional details in Text S2).

### Interpretation using binary outcomes (seroprevalence)

Many current rapid diagnostics for neglected tropical diseases and malaria provide dichotomous (seropositive/seronegative) results [31,32]. The nonparametric method is very general and can, in principle, be applied to dichotomous test results, where *E*(*Y*_*a,x*_) then estimates the age-dependent seroprevalence curve and *E*(*Y*_*x*_) estimates age-adjusted mean seroprevalence. We compared quantitative antibody curves with seroprevalence curves for the Mauke lymphatic filariasis and Garki malaria examples where we could establish clear seropositivity cutoffs. In the Mauke study, a previous analysis found a cutoff value of 10968 light units had sensitivity >98% and specificity ranging from 94-100% when compared with negative and positive control samples, including those co-infected with other helminths [16]. In the Garki study, we treated any response to *P. falciparum* in the IFA test as positive [17]. For enteric pathogens, we used finite mixture models to estimate 2 gaussian components [33], and then estimated the seropositivity cutoff values using the mean plus 3 standard deviations of the first component; we compared three separate cutoffs using USA samples alone, Haiti samples alone, and combined USA and Haiti samples.

### Software

We used the R statistical software (version 3.2.4, www.R-project.org) for analysis and data visualization. Full replication files (datasets, scripts) and an R software package to easily implement the approach (tmleAb) are available through GitHub and the Open Science Framework: https://osf.io/8tqu4.

### Ethics statement

Study protocols for Mauke were approved by the government of the Cook Islands and the NIAID Institutional Review Board, and all adult subjects provided written informed consent. Consent for children was obtained by verbal assent as well as written consent from legal guardians. The Haiti study protocol was reviewed and approved by the Centers for Disease Control and Prevention’s Institutional Review Board and the Ethical Committee of St. Croix Hospital (Leogane, Haiti) and all subjects provided verbal consent. Human subjects review boards approved a verbal consent process because the study communities had low literacy rates. Mothers provided consent for young children, and children 7 years or older provided assent.

## Results

### Lymphatic filariasis in Mauke

There was a distinct shift in the *W. bancrofti* Wb123 age-antibody curve before and five years after a single diethylcarbamazine MDA (Figure 1a), and differences between curves show more gradual antibody acquisition with age in the post-MDA measurement (Figure 1b). As previously noted [15], mean Wb123 antibody levels declined in individuals who tested positive for circulating filarial antigen before MDA (a sign of active infection) but had no detectable circulating antigen post-MDA, as well as among those who tested negative for circulating antigen at both time points (Figure 1c). Together, these results show that slower antibody acquisition combined with antibody loss, presumably a reflection of lowered transmission potential post-MDA, underlie the curve shift. Seroprevalence estimates for Wb123 followed a similar pattern as the quantitative antibody response (Figure S1). A caveat of the Wb123 seroprevalence analysis was that the seropositivity cutoff, chosen to have near perfect sensitivity and specificity with respect to controls [16], fell in the center of the Wb123 distribution in the post-MDA measurement (lower transmission) (Figure S1). This observation makes it more difficult to argue that there were two distinct seropositive and seronegative populations -- an assumption avoided when relying directly on quantitative antibody levels.

**Figure 1:**
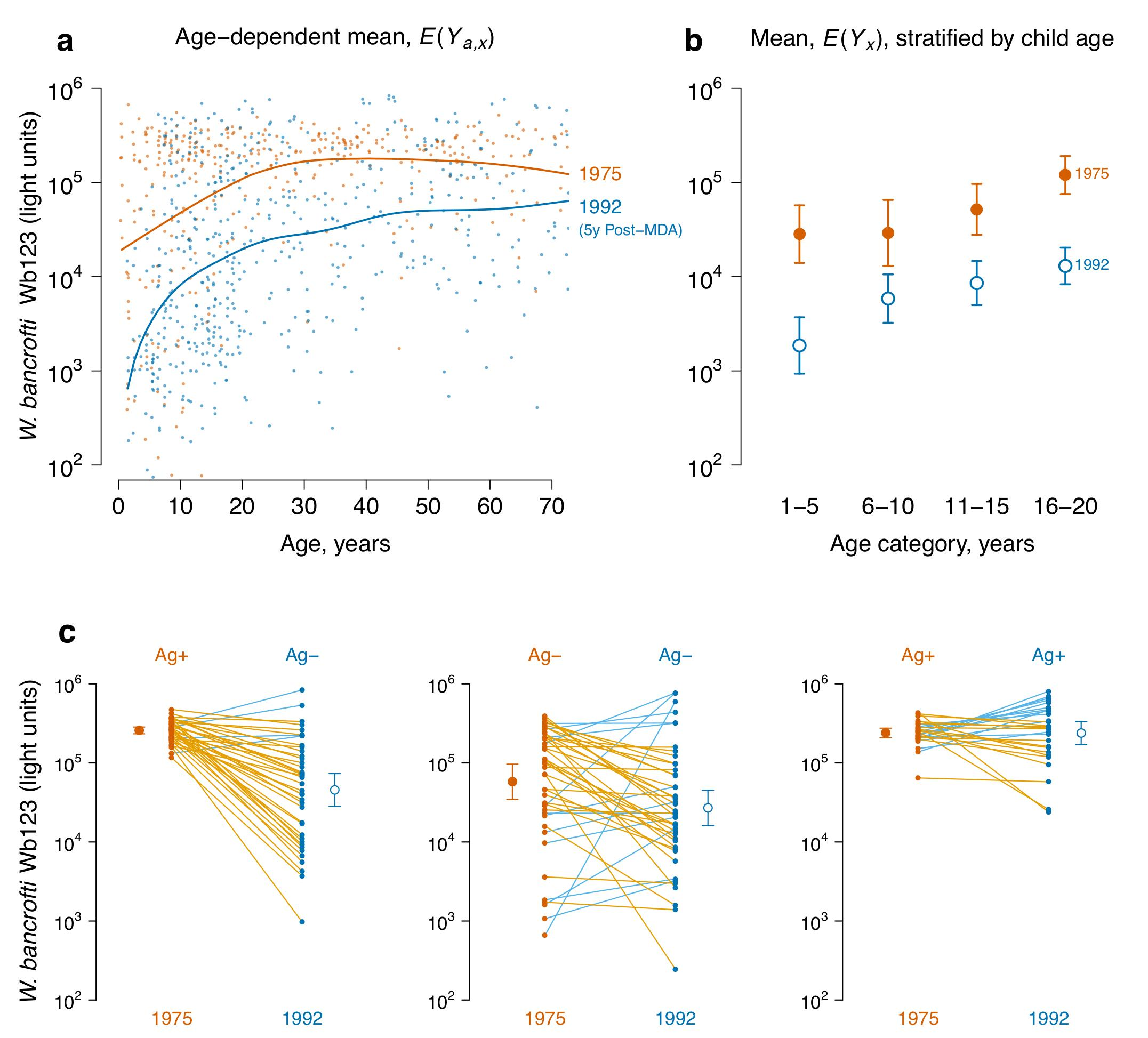
A shift in the *Wuchereria bancrofti* Wb123 age-antibody curve measures a reduction in transmission due to mass drug administration (MDA) on Mauke Island. IgG antibody response to the Wb123 antigen for *W. bancrofti* measured in blood specimens from residents in 1975 (N=362) before MDA and again in 1992 (N=553), five years following a single, island-wide MDA with diethylcarbamazine. **a,** Mean antibody levels *E*(*Y*_*a,x*_) by age (*a*) and survey year (*x*); individual antibody responses (points) are shown along with summary curves. **b,** Age-adjusted geometric mean antibody response, *E*(*Y*_*x*_), and 95% confidence intervals before (1975) and five years after (1992) MDA, stratified by 5 year age category (all differences significant at P ≤ 0.0001 after Bonferroni correction). **c,** Wb123 antibody response in 1975 and 1992 stratified by the presence of circulating filarial antigens (Ag) at each measurement in the subsample of 112 individuals who were measured at both time points (two individuals not shown were Ag- in 1975 and Ag+ in 1992), along with age-adjusted geometric means, *E*(*Y*_*x*_), and 95% confidence intervals. Differences between means are significant (Bonferroni corrected *P* ≤ 0.01) except for the Ag+/Ag+ group. Individual trajectories are colored by the higher of the two measurements: decreases are or- ange, increases are blue. The source data used to generate this figure are here: https://osf.io/8tqu4 (mauke), and the scripts used to generate the figure are here: https://osf.io/ek3sx (mauke).

### Malaria in the Garki Project, Nigeria

Compared to control villages, there was a consistent shift in *P. falciparum* age-antibody curves with increased length of the insecticide spraying and MDA intervention in the Garki project (Figure 2a). During the active intervention period, children in intervention villages exhibited a sharp drop in antibody levels from birth and a more gradual increase in antibody levels compared with children in control villages. Mean IFA titers demonstrated group comparability before intervention, reduced transmission during intervention, and a transmission resurgence after the intervention period (Figure 2a) -- a pattern that corresponded closely with rates of patent parasitemia measured in the original study [17]. Age-dependent seroprevalence curves followed a similar pattern to the quantitative antibody results, but changes due to intervention were less pronounced because reductions in seroprevalence were only detectable among children < 5 years old (Figure 2b). When estimated at the finer resolution, village level rather than in aggregate, mean antibody titers more clearly distinguished intervention and control villages compared with seroprevalence (Figure S2). Village level mean antibody titers correlated strongly with wet season EIR (Spearman’s ρ=0.75) and with seroprevalence (Spearman’s ρ=0.93, Figure 3).

**Figure 2:**
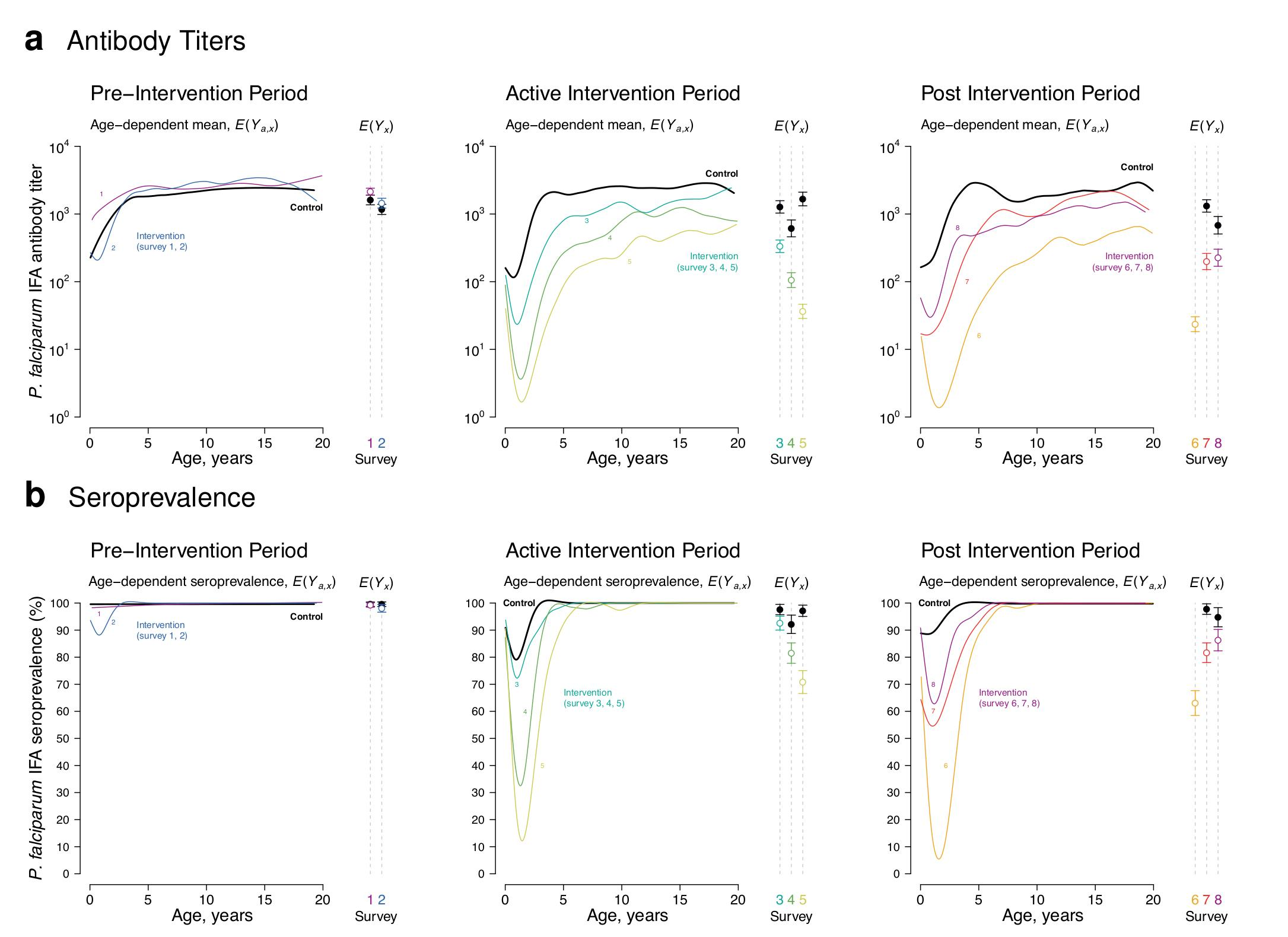
Shifts in the *Plasmodium falciparum* age-antibody curve measure changes in malaria transmission due to intervention in the Garki Project, Nigeria (1970-1976). Antibody response measured with the IgG indirect fluorescent antibody (IFA) test for P. falciparum using semi-quantitative antibody titers (**a**) or reduced to seroprevalence (**b**). Estimates stratified by pre-intervention period wet and dry seasons (survey rounds 1-2), active intervention period (survey rounds 3-5, at 20, 50, and 70 weeks following the start of intervention), and the post-intervention period (survey rounds 6-8 at 20, 40, and 90 weeks following the end of the intervention). N=4,774 total measurements, with 153 - 442 measurements per curve. Control measurements were combined across survey rounds within each period when plotting the curves to facilitate visual comparison of shifts in transmission between surveys. Age-adjusted means by intervention group, *E*(*Y_x_*), provide summary differences between curves at each survey round. Error bars show 95% confidence intervals for the age-adjusted geometric means (**a**) or seroprevalence (**b**) and differences between groups are significant *P* ≤ 0.01 (Bonferroni corrected) for all rounds except pre-intervention surveys 1 and 2. Control villages were not measured in survey 6. The source data used to generate this figure are here: https://osf.io/8tqu4 (garki), and the scripts used to generate the figure are here: https://osf.io/ek3sx (garki).

**Figure 3:**
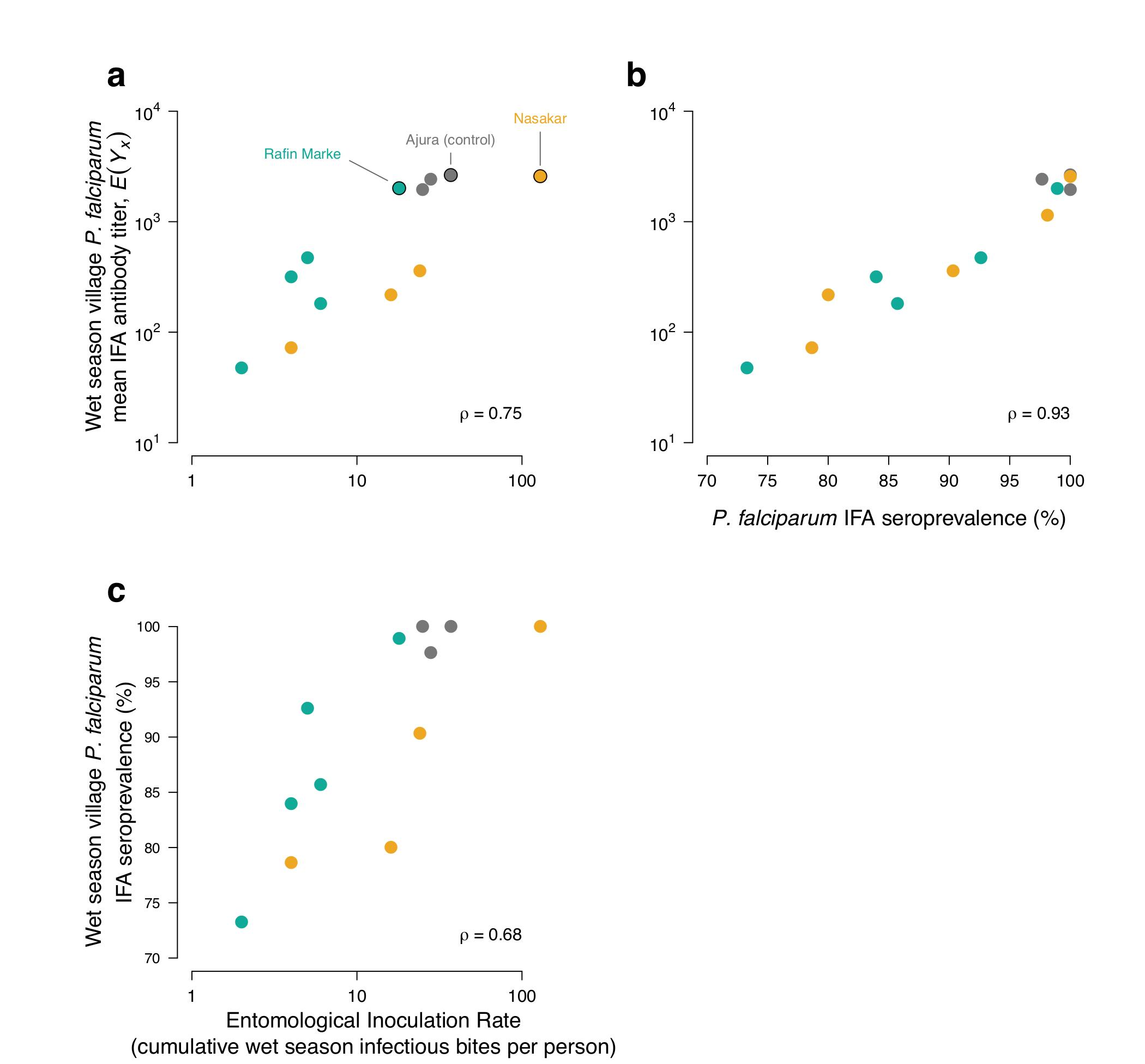
Comparison of mean *Plasmodium falciparum* IFA antibody titers with wet season entomological inoculation rate (EIR) and IFA seroprevalence in the three study villages with paired entomological and serological measurements. **a,** *P. falciparum* IFA titers versus EIR. **b** *P. falciparum* IFA titers versus seroprevalence. **c,** *P. falciparum* seroprevalence versus EIR. Ajura was a control village (no treatment) while Rafine Marke and Nasakar were intervention villages. A single data point outside the figure range is not shown in EIR plots (Nasakar 1972, EIR value = 0, *E* (*Y*_*x*_) = 103.0591), but was included in the Spearman’s rank correlation estimates (*ρ*). The source data used to generate this figure are here: https://osf.io/8tqu4 (garki), and the scripts used to generate the figure are here: https://osf.io/ek3sx (garki).

Malaria transmission was highly seasonal during the study, with more intense vector transmission and incident infections in the wet seasons [17]. In a secondary analysis, we restricted the population to control villages and fit age-antibody curves separately by survey rounds 1-5, which corresponded to sequential wet and dry seasons. We observed a distinct shift in the age-antibody curve, consistent with lower transmission in the dry season, but only among children <5 years old; older children exhibited far less seasonal variation in mean IFA antibody titers compared with children <5 years (Figure 4).

**Figure 4:**
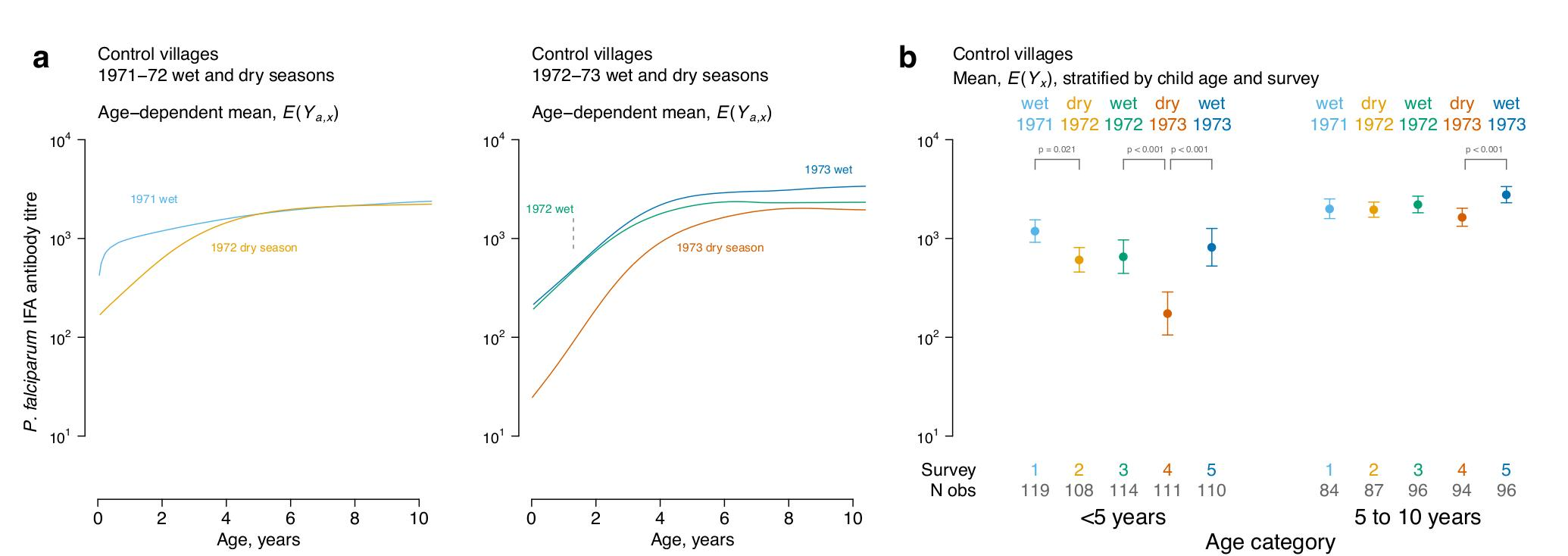
Higher sensitivity among children <5 y to seasonal changes in *Plasmodium falciparum* transmission as depicted by age-antibody curves estimated within control villages in the Garki Project, Nigeria (1970-1976). Antibody response measured with the IgG indirect fluorescent antibody (IFA) test for *P. falciparum*. **a,** Mean antibody levels by age (***a***) and season (*x*), *E*(*Y*_*a,x*_). **b,** Age-adjusted geometric means by age category and season, *E*(*Y*_*x*_), summarize the curves. Error bars show 95% confidence intervals and *P*-values mark significant differences (Bonferroni corrected) between adjacent seasons. The source data used to generate this figure are here: https://osf.io/8tqu4 (garki), and the scripts used to generate the figure are here: https://osf.io/ek3sx (garki).

### Enteric pathogens in Haiti and the USA

Age-antibody curves for IgG antibody responses to protozoan, bacterial, and viral enteric pathogens were consistent with lower levels of enteric pathogen transmission in the USA (Figure 5). The Haiti and USA populations likely illustrate enteric antibody curves near the bounds of high and low transmission environments, and show that as transmission declines the curves flatten. The results illustrate both the consistency of the general pattern across diverse taxa as well as the facility with which the analysis method generalizes to multiplex applications where numerous antibodies can be measured from a single blood spot. In most cases, enteric pathogen antibody distributions did not show obvious seropositive and seronegative subpopulations, and seropositivity cutoff values varied when estimated using different sample sets (Figure S3). In most cases, seropositivity cutoffs using the Haiti specimens alone fell outside the observed range of the antibody distributions (Figure S3).

**Figure 5.**
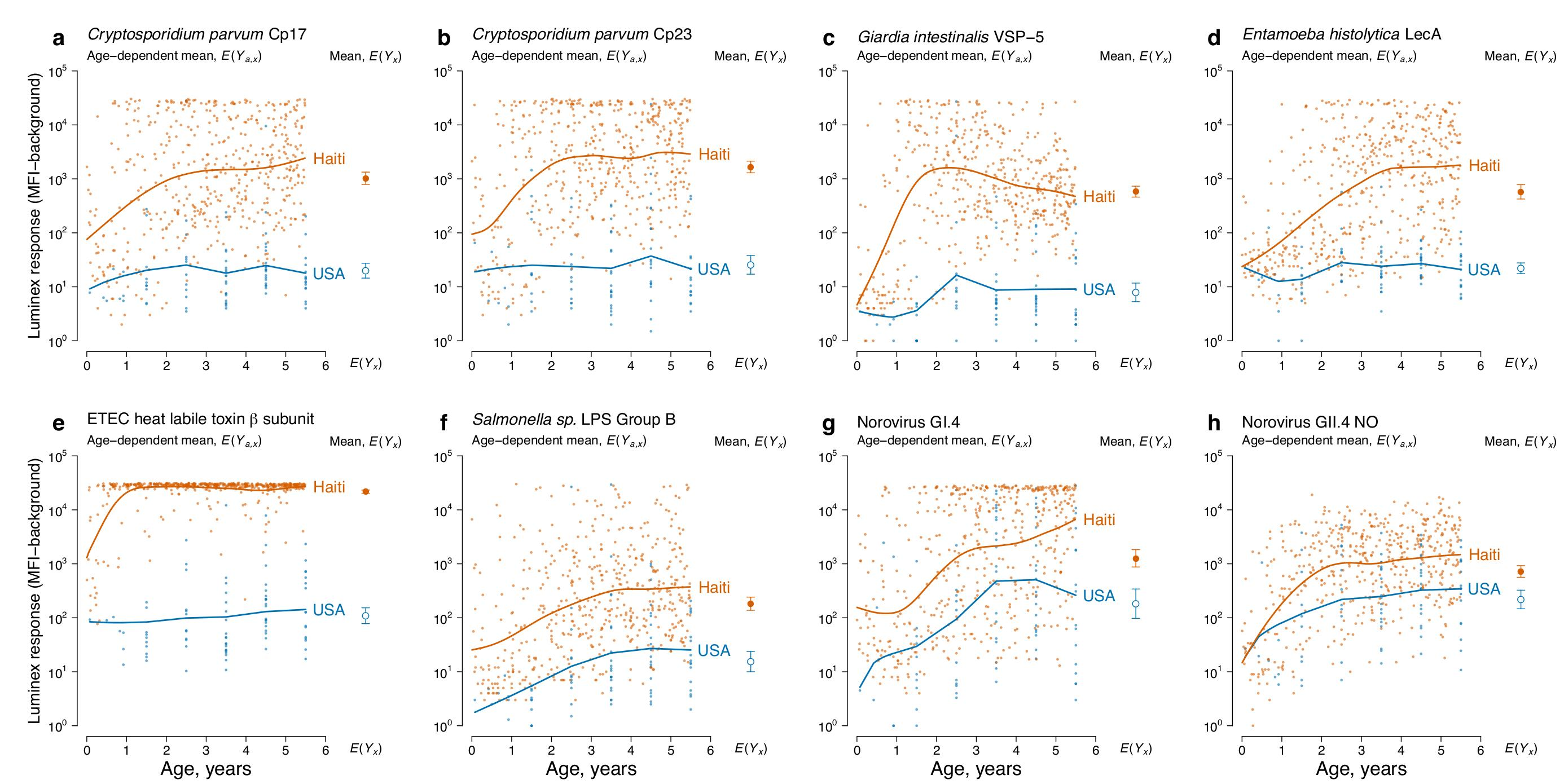
Differences in enteric pathogen transmission between children in Leogane, Haiti (N=511) and the United States (USA) (N=86) measured by age-antibody curves. Antibody response measured as median fluorescence intensity (MFI) minus background in multiplex bead assays on the Luminex platform. In each panel, individual antibody responses (points) are shown along with age-dependent means. Each panel also includes the geometric mean by country, *E*(*Y*_*x*_), with error bars marking 95% confidence intervals (all differences significant at *P* ≤ 0.001 after Bonferroni correction). **a.** *Cryptosporidium parvum* recombinant 17-kDa antigen; **b.** *Cryptosporidium parvum* recombinant 27-kDa antigen; **c.** *Giardia intestinalis* variant-specific surface protein-5 (VSP-5); **d.** *Entamoeba histolytica* lectin adhesion molecule (LecA); **e.** enterotoxigenic *Escherichia coli* (ETEC) heat labile toxin *β* subunit; **f.** *Salmonella spp.* lipopolysaccharide (LPS) Group B; **g.** Norovirus Group I.4; **h.** Norovirus Group II.4 New Orleans. The source data used to generate this figure are here: https://osf.io/8tqu4 (enterics), and the scripts used to generate the figure are here: https://osf.io/ek3sx (enterics).

## Discussion

### Key findings

We have shown that diverse, pathogen-specific serum IgG levels follow a characteristic shape with increasing age, and that changes in transmission are reflected in a shift of the age-antibody curve that can be summarized by changes in mean antibody levels. Consistent with our hypothesis, reduced transmission produced age-antibody curves that rose more slowly and plateaued at lower levels. The generality and consistency of the age-antibody relationship across diverse infectious diseases, populations, and diagnostic platforms suggest that this simple, robust methodology constitutes a useful way to measure changes in transmission for pathogens with serum IgG antigen targets.

### Interpretation

Our results support at least three reasons to consider the use quantitative antibody levels to measure changes in pathogen transmission as a complement or alternative to seroprevalence and other metrics based on a binary response. First, analyses of mean antibody levels avoid the need to define seropositivity cutoffs and readily accommodate pathogens where antibody levels wane with time since exposure. The *W. bancrofti* and enteric pathogen analyses provided many examples where seropositivity cutoffs could either not be estimated or fell in the center of symmetric distributions (Figures S1, S3). In those cases, it was difficult to assume distinct seropositive and seronegative populations, whereas a comparison based on mean antibody levels avoided that assumption. Second, in settings where most individuals fall above or below the seropositivity cutoff, analyses of mean antibody levels can provide a higher resolution measure of changes in transmission compared with seroprevalence. In the Garki project, *P. falciparum* transmission reductions evident across ages 0-20 years with quantitative antibodies were only detectable among children <5 years with seroprevalence (Figure 2). Waning *W. bancrofti* Wb123 antibody levels among individuals in Mauke without circulating antigen (Figure 1c) further underscored how quantitative responses could provide more information about gradations in exposure that are lost with binary, positive/negative assays. These findings are broadly consistent with recent comparisons of estimates from quantitative antibody and seroprevalence estimates in the malaria context [7]. Indeed, quantitative antibody levels could provide complementary, high resolution information alongside more traditional metrics of infection to identify heterogeneous transmission in populations -- a recent example illustrated the value of using malarial antibody levels directly to identify transmission hotspots in Cambodia [34], and similar applications could be possible for NTDs and other infectious diseases. A third consideration is that mean antibody levels should require fewer observations to estimate precisely than seroprevalence since reducing a quantitative measure to a binary measure results in a theoretical loss of >36% of Fisher’s information [35]. A sample of 20 individuals is unlikely to provide accurate information about seroprevalence or seroconversion rates [6], but could provide a reliable estimate of mean antibody levels -- the village-level analyses in the Garki project showed that use of *P. falciparum* quantitative antibodies led to larger and more precise estimates of differences between control and intervention groups than seroprevalence when estimated in small samples (Figure S2). This could be a particular advantage for serological surveillance in population-based surveys where sampling clusters often include fewer than 30 people [36], and our labs are currently working on more formal guidance for sampling designs based on quantitative antibody levels.

The use of data-adaptive, ensemble machine learning to fit antibody curves and compare means has several strengths in the context of developing a generalized methodology for integrated surveillance. The approach is: extremely flexible, easy to implement with open-source software (replication files include an R package and vignette: https://osf.io/8tqu4), easy to adjust for potential confounding covariates, minimally biased, and highly efficient [14,29,37]. Ensemble approaches have been very successful in applications where no single model is likely to be correct across diverse applications -- for example, cause of death classification in the Global Burden of Disease studies [38], mortality prediction in intensive care units [39], or predicting malaria incidence from diverse antibody panels [40]. The ensemble library can include a range of models or algorithms, and if simpler models perform better they will be upweighted in the estimation [14]. Previous statistical methods have used quantitative antibody levels to measure differences in pathogen transmission by estimating parameters such as infection rates [40–42], seroconversion rates [3,4,43], or antibody acquisition rates [7,8,43]. Incidence and seroconversion rates are epidemiologically useful, but to estimate them from quantitative antibody levels requires strong modeling assumptions, or well-characterized longitudinal cohorts that directly measure the parameter of interest to train models, or both. Measuring differences in transmission directly from antibody levels with age-antibody curves requires neither modeling assumptions nor well-characterized cohorts to train models or fit parameters. This could be an advantage for integrated surveillance platforms where pathogens vary greatly in their specific immunology and no detailed longitudinal cohorts with antibody infection profiles exist for most pathogens. The ensemble fits revealed consistent shifts in the age-antibody curve with lower transmission, but individual curves followed complex age-dependent patterns that varied by pathogen and setting. Data-adaptive, nonparametric algorithms tended to perform better than simpler models in terms of cross-validated *R*^*2*^, but there was no member of the ensemble that performed best across all pathogens and transmission settings (Figure S4). We included in the ensemble library an antibody acquisition model developed for malaria [7], but that particular model underperformed in comparison with more flexible algorithms such as smoothing splines (Figure S4). This result suggests it may be difficult to devise a single model that describes the full diversity of age-dependent antibody response across very different infectious diseases, and underscores the value of considering an ensemble approach for broad analyses envisioned through integrated surveillance.

The specific antibody kinetics and the age range in which the curves are estimated will influence the sensitivity of this approach to detect changes in transmission. Curves fit using antibodies with shorter half-lives should theoretically exhibit shifts more quickly with changes in transmission. Microarray screening efforts to identify malarial antibodies with a range of half-lives [40] open the possibility for discovering antibodies with high sensitivity to measure changes in transmission over short periods. With antibodies measured in multiplex, future work could develop methods to combine multiple antigens expressed by the same pathogen into a single quantitative response -- a composite measure could prove more robust to differential immunogenicity arising from differences in host genetics.

Our analyses show that serological surveillance among children captures the period of greatest change in the age-antibody curve, and analyses using children would be less susceptible to longer-term “cohort effects” that could influence the age-antibody relationship for antibodies with long half-lives [44]. Children are likely the most sensitive population to measure reductions in transmission: age-specific immunological profiles of malaria and vaccine response to diverse pathogens show that young children lose antibodies more quickly than adults because short-lived B cells predominate in young children, and antigen presentation and helper T-cell function increase with age [45–47]. Surveillance activities that measure a very narrow age range, such as transmission assessment surveys to monitor lymphatic filariasis elimination programs (which only measure children ages 6-7 years), cannot estimate a full age-antibody curve but the summary mean would still provide a robust measure of adjusted mean antibody levels to compare populations (Fig 1b).

Quantitative IgG antibody response integrates information about an individual’s pathogen exposure over time [3] -- a characteristic of particular import for community-based surveillance of pathogens with low annual incidence and pathogens that cause many asymptomatic infections. Low incidence and asymptomatic presentation make community-based surveillance of changes in transmission difficult because either scenario requires very large numbers of specimens to be tested to identify incident infections. For example, *Cryptosporidium parvum* is implicated as a major pathogen of concern due to its contribution to hospitalized cases and prolonged episodes of diarrhea [48], but community-based studies of *Cryptosporidium sp.* require the collection of thousands of stool specimens. Large studies are needed because, even in hyper-endemic settings, rates of incident infections fall below a single episode per person-year [49], and because intermittent shedding of small numbers of oocysts in the stools of some infected individuals can make detection difficult [50]. We have illustrated that full age-antibody curves can be estimated with as few as 100 observations spread over different ages, which suggests they could be useful in the surveillance of pathogens with low annual incidence, or asymptomatic infections that clinical surveillance activities typically miss.

### Limitations and next steps

There are two main limitations of the approach. First, mean antibody levels do not estimate a direct epidemiologic transmission parameter, such as the incidence or force of infection. Thus, while mean antibody levels provide a flexible, sensitive method to measure differences in transmission within- or between populations, they provide only indirect information about the relative importance or health burden of different pathogens. Using the same underlying statistical method with binary outcomes to estimate seroprevalence (Figures 2, S1) partly addresses this limitation at the cost of losing some information, and our labs are actively working to extend these general methods to estimate a pathogen’s force of infection. A second limitation is that if a quantitative antibody assay has no global reference standard to translate arbitrary units into antibody titers, it will be difficult to make direct comparisons of mean antibody levels across different assays and studies. Assay standardization is a common challenge of any serological surveillance, so this limitation is shared by all methods that measure changes in transmission from antibody assays. The development of global reference standards for antibody assays used in infectious disease surveillance [51], as currently exist for many vaccine-preventable diseases, would facilitate between-study comparisons.

There are two main limitations of the approach. First, mean antibody levels do not estimate a direct epidemiologic transmission parameter, such as the incidence or force of infection. Thus, while mean antibody levels provide a flexible, sensitive method to measure differences in transmission within- or between populations, they provide only indirect information about the relative importance or health burden of different pathogens. Using the same underlying statistical method with binary outcomes to estimate seroprevalence (Figures 2, S1) partly addresses this limitation at the cost of losing some information, and our labs are actively working to extend these general methods to estimate a pathogen’s force of infection. A second limitation is that if a quantitative antibody assay has no global reference standard to translate arbitrary units into antibody titers, it will be difficult to make direct comparisons of mean antibody levels across different assays and studies. Assay standardization is a common challenge of any serological surveillance, so this limitation is shared by all methods that measure changes in transmission from antibody assays. The development of global reference standards for antibody assays used in infectious disease surveillance [51], as currently exist for many vaccine-preventable diseases, would facilitate between-study comparisons.

There are two main limitations of the approach. First, mean antibody levels do not estimate a direct epidemiologic transmission parameter, such as the incidence or force of infection. Thus, while mean antibody levels provide a flexible, sensitive method to measure differences in transmission within- or between populations, they provide only indirect information about the relative importance or health burden of different pathogens. Using the same underlying statistical method with binary outcomes to estimate seroprevalence (Figures 2, S1) partly addresses this limitation at the cost of losing some information, and our labs areactively working to extend these general methods to estimate a pathogen’s force of infection. A second limitation is that if a quantitative antibody assay has no global reference standard to translate arbitrary units into antibody titers, it will be difficult to make direct comparisons of mean antibody levels across different assays and studies. Assay standardization is a common challenge of any serological surveillance, so this limitation is shared by all methods that measure changes in transmission from antibody assays. The development of global reference standards for antibody assays used in infectious disease surveillance [51], as currently exist for many vaccine-preventable diseases, would facilitate between-study comparisons.

This study focused on IgG responses to lymphatic filariasis, malaria, and enteric pathogens measured in blood, but the method should apply to other immunoglobulin isotypes, other specimen types, and other infectious diseases. For example, similar shifts in IgE curves have been documented in populations with different soil transmitted helminth transmission [11], salivary IgG and IgA norovirus assays have been developed [52], and NTDs such as trachoma [53], dengue [54], and chikungunya [55] all have well-defined antigens that would be amenable to this methodology. Mean antibody response in defined geographic areas over time could translate directly to mapping activities used to target intervention programs and monitor transmission or immunization coverage. The ability to combine dozens of recombinant antigens into multiplex bead assays opens the possibility for high-throughput, integrated infectious disease surveillance that includes pathogens targeted for elimination such as NTDs and malaria alongside newly emerging pathogens, and vaccine preventable diseases [51]. The methods developed here provide a very general tool for integrated surveillance of antibody response from such data.

## Acknowledgements

We thank Kunal Mishra at the University of California, Berkeley for assistance with software package development (tmleAb), and J. Vinje and V. Costantini at the CDC for gifts of reagents.

---

**Figure S1:**
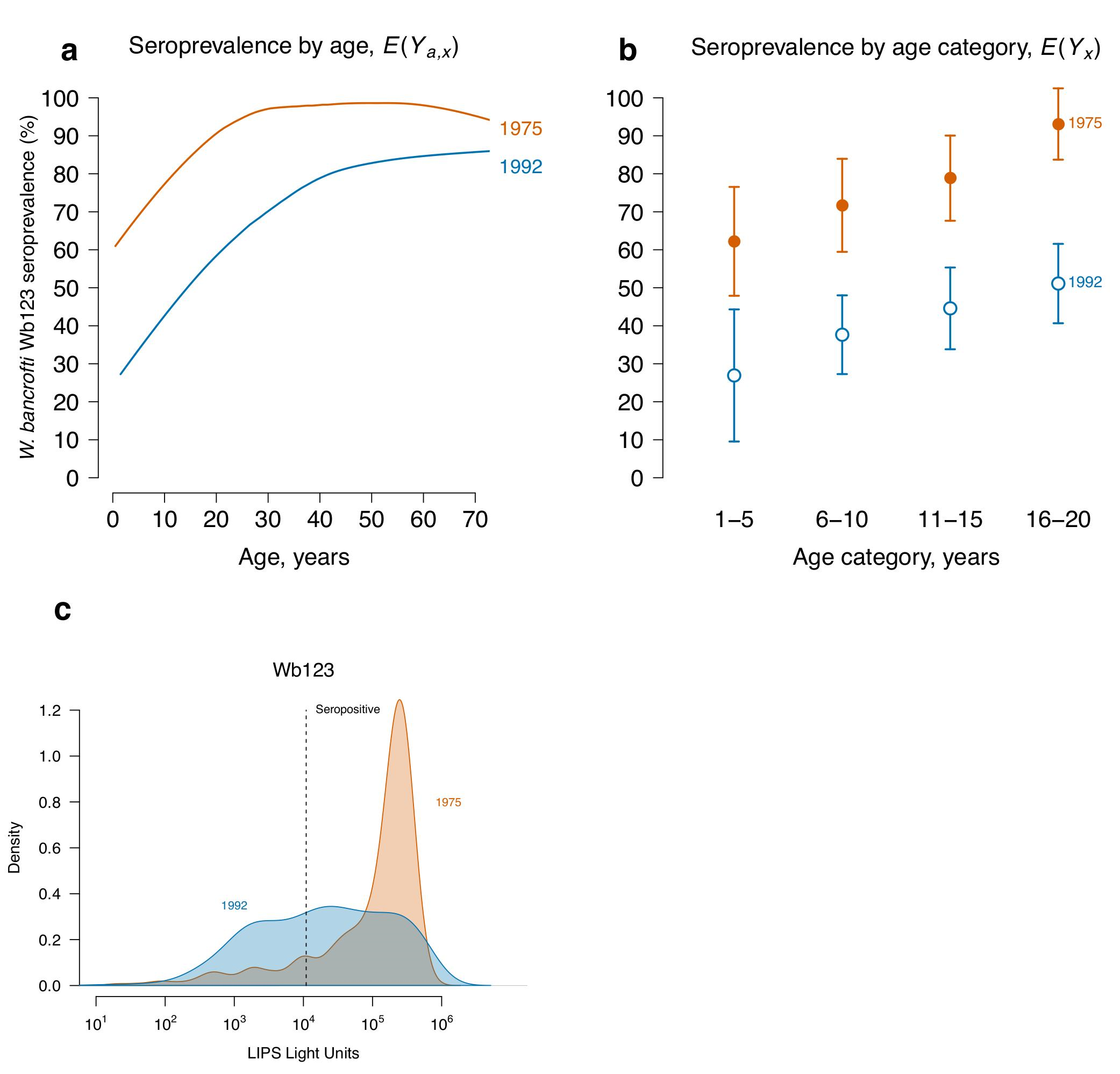
A shift in the *Wuchereria bancrofti* Wb123 age-dependent seroprevalence curve measures a reduction in transmission due to mass drug administration (MDA) on Mauke Island. **a**, *W. bancrofti* Wb123 age-dependent seroprevalence curves. **b**, mean Wb123 seroprevalence by age category. Wb123 measured from blood specimens collected from residents in 1975 (N=362) before MDA and again in 1992 (N=553), five years following a single, island-wide MDA with diethylcarbamazine. **c**, Kernel smoothed density distributions of Wb123 antibody levels in 1975 and 1992, along with a seropositivity cutoff value (10968 light units) identified to maximize sensitivity and specificity using positive and negative controls (Kubofcik et al. PLOS Negl Trop Dis. 2012;6: e1930) and used in the seroprevalence analyses. The source data used to generate this figure are here: https://osf.io/8tqu4 (mauke), and the scripts used to generate the figure are here: https://osf.io/ek3sx (mauke).

**Figure S2:**
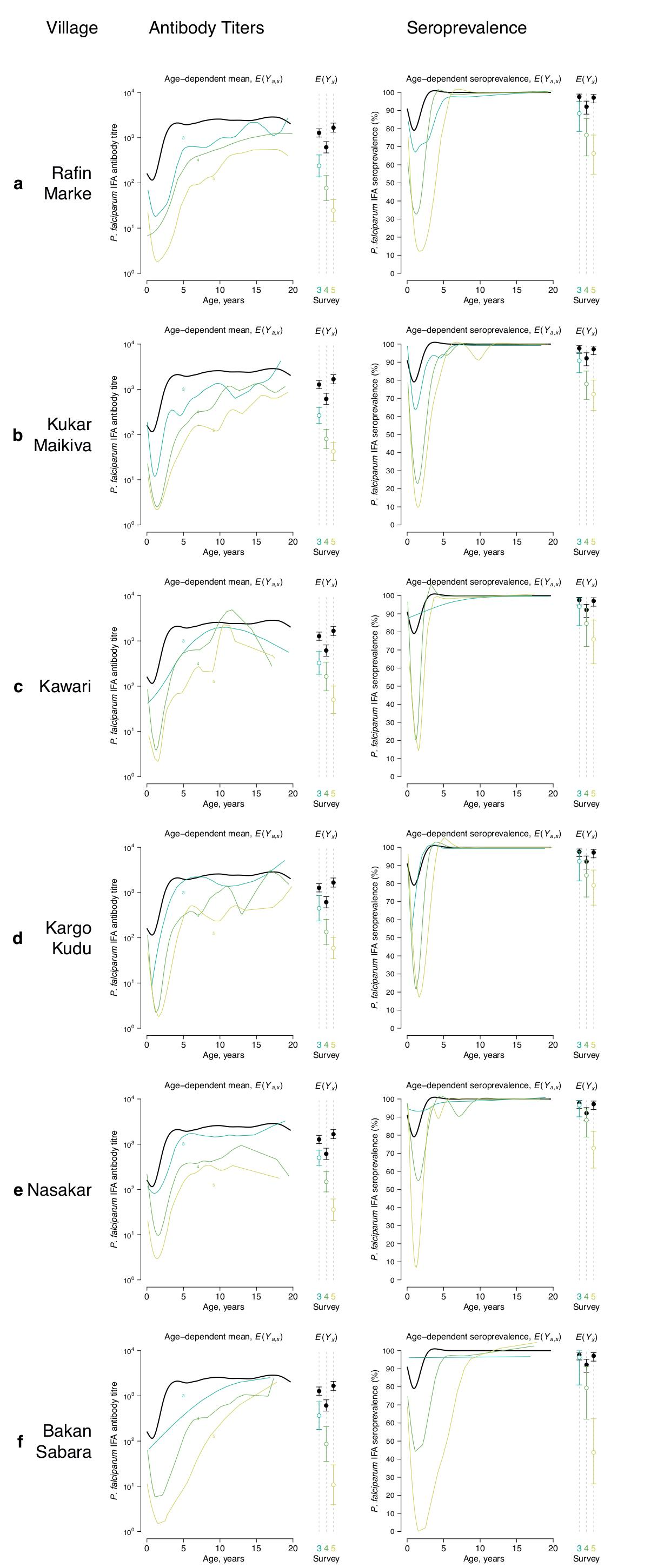
*Plasmodium falciparum* age-antibody curves estimated separately for each intervention village (**a-f**) measure reductions in malaria transmission due to intervention in the Garki Project, Nigeria (1970-1976). Antibody response measured with the indirect fluorescent antibody (IFA) test for *P. falciparum* during the active intervention period (survey rounds 3-5, at 20, 50, and 70 weeks following the start of intervention). The left column includes curves and means using quantitative antibody titers and the right column includes estimates using seroprevalence. Each curve and mean was estimated with between 19 and 119 observations, depending on the village and survey round. Error bars show 95% confidence intervals for the summary means. Each village panel includes the same curve for the two control villages (black) for comparison. Control measurements were combined across survey rounds within each period when plotting the curves to facilitate visual comparison of shifts in transmission between surveys. The source data used to generate this figure are here: https://osf.io/8tqu4 (garki), and the scripts used to generate the figure are here: https://osf.io/ek3sx (garki).

**Figure S3:**
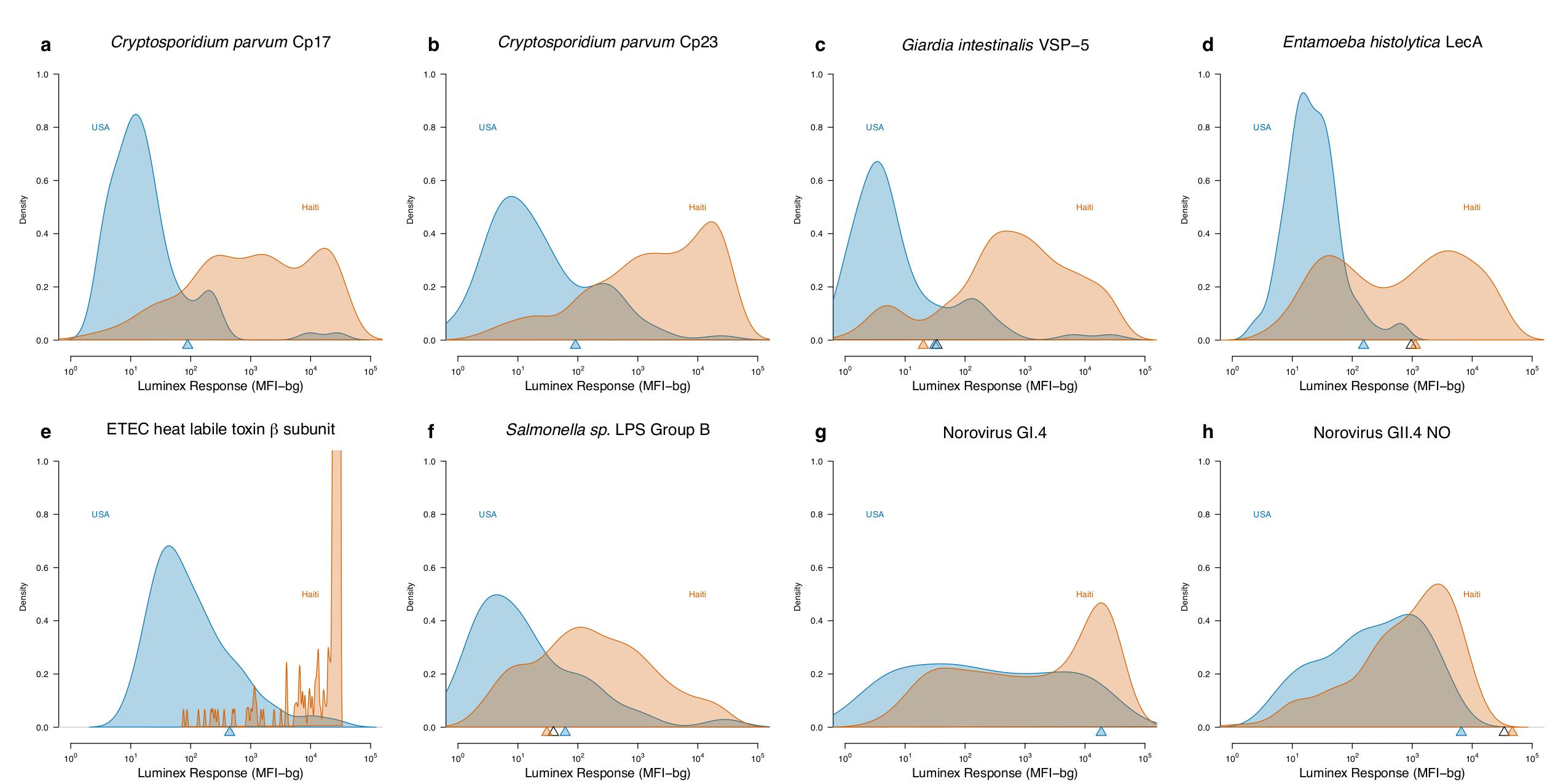
Kernel density smoothed distributions of enteric antibody response in children < 5.5 years old in the United States (USA) and Haiti. Antibody response measured in multiplex on the Luminex platform using median fluorescence intensity minus background (MFI-bg). Triangles below the distributions indicate seropositivity cutoffs determined using finite gaussian mixture models fit to each country’s measurements (filled triangles) or the combined sample set (open triangle), where cutoffs were determined using the mean+3*SD of the first gaussian component. Cutoff values beyond the range of observed measurements are not shown. **a.** *Cryptosporidium parvum* recombinant 17-kDa antigen; **b.** *Cryptosporidium parvum* recombinant 27-kDa antigen; **c.** *Giardia intestinalis* variant-specific surface protein-5 (VSP-5); **d.** *Entamoeba histolytica* lectin adhesion molecule (LecA); **e.** enterotoxigenic *Escherichia coli* (ETEC) heat labile toxin *β* subunit. Note: the Y-axis is truncated at 1.0 but extends to 10.0 for this antibody; **f.** *Salmonella spp.* lipopolysaccharide (LPS) Group B; **g.** Norovirus Group I.4; **h.** Norovirus Group II.4 New Orleans. The source data used to generate this figure are here: https://osf.io/8tqu4 (enterics), and the scripts used to generate the figure are here: https://osf.io/ek3sx (enterics).

**Figure S4:**
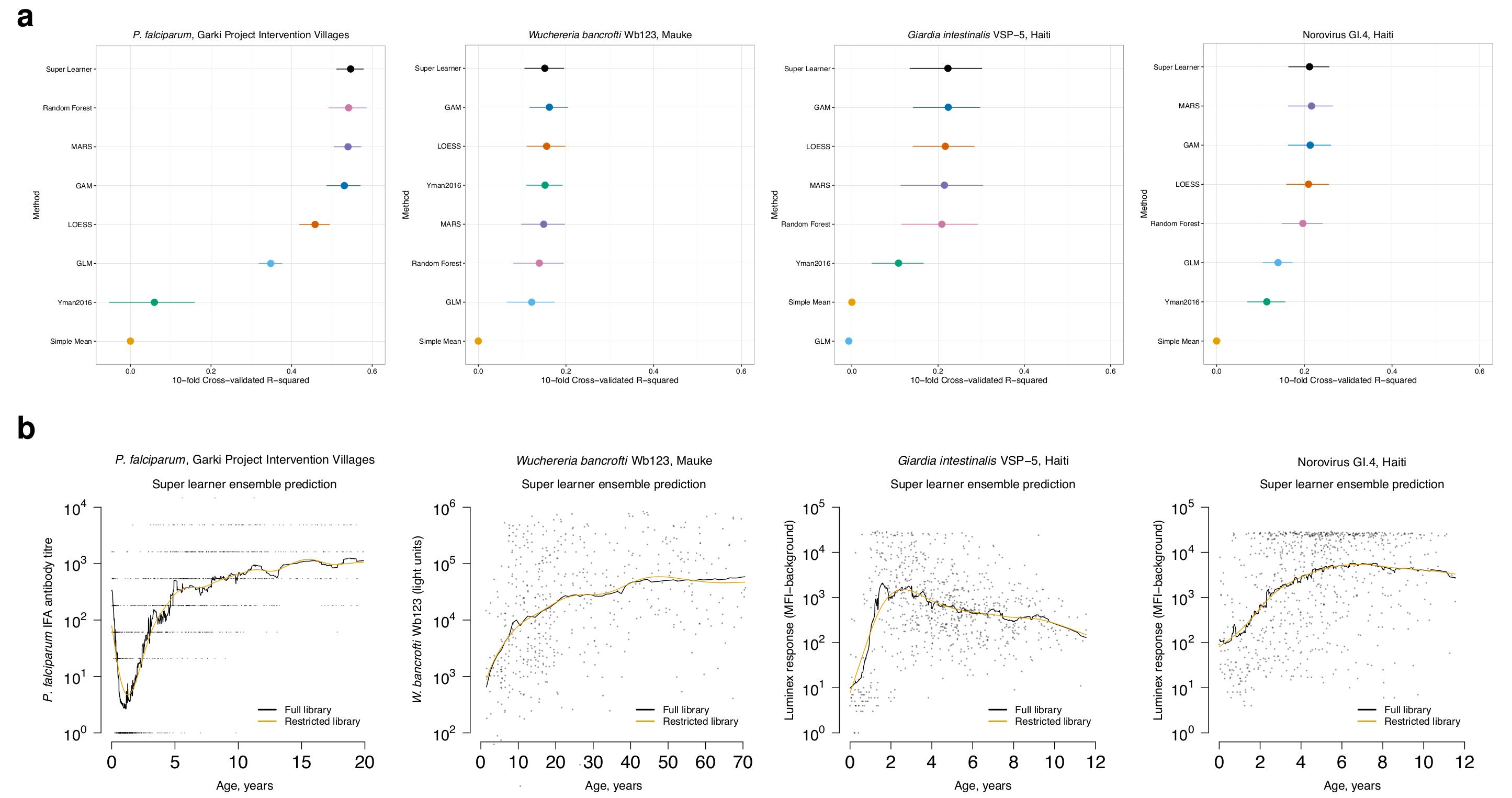
Performance of the ensemble and individual models/algorithms. The super learner algorithm combines predictions from an ensemble by selecting a weighted combination of individual model predictions using cross-validation to optimize the bias-variance tradeoff – in this case, mean squared error (MSE). We estimated the cross-validated MSE and the related cross-validated R2 of the super learner along with each of its constituent models/algorithms across a range of populations and pathogens in the analysis. Cross-validated *R*^2^ represents the percentage of outcome variability explained beyond estimating the simple mean. **a**, Cross-validated estimates ofR2for the super learner ensemble and its constituent models/algorithms across example populations and pathogens. Horizontal lines represent twice the standard error of the *R*^2^ estimates measured across 10 cross-validation splits. In all cases, the algorithms only included age as a feature in antibody level prediction. In the Garki Project, where additional information was available, including additional covariates (sex, wet vs. dry season, village membership) did not markedly improve *R*^2^ for any algorithm or the ensemble. **b** Super learner ensemble estimates of age-dependent antibody curves for different populations and pathogens including the full library (all members listed in **a**) as well as a restricted library that excluded two highly adaptive algorithms (Random Forest and MARS).The most highly adaptive algorithms that we considered (random forest and MARS) often led to jagged age-antibody curves, and excluding them led to consistent but smoother curves. We therefore used the restricted library in other analyses. Abbreviations: GAM: generalized additive models with natural splines; GLM: generalized linear model; LOESS: Locally weighted regression; MARS: Multivariate adaptive regression splines; Yman2016: Antibody acquisition model proposed by Yman et al. [Sci Rep 2016; 6:19472]. The source data used to generate this figure are here: https://osf.io/8tqu4, and the scripts used to generate the figure are here: https://osf.io/ek3sx.

## Supporting Information Captions

**Text S1** Relationship between the age-adjusted mean antibody response and the area under the curve

**Text S2** Technical details of estimating age-dependent antibody curves and changes in mean antibody levels

**Table S1** STROBE checklist

**Figure S1** A shift in the *Wuchereria bancrofti* Wb123 age-dependent seroprevalence curve measures a reduction in transmission due to mass drug administration on Mauke Island.

**Figure S2** *Plasmodium falciparum* age-antibody curves estimated separately for each intervention village

**Figure S3** Kernel density smoothed distributions of enteric antibody response in children < 5.5 years old in the United States and Haiti

**Figure S4** Performance of the ensemble and individual models/algorithms.

## Funding Statement

This work was supported by National Institute of Allergy and Infectious Diseases grants K01AI119180 (BFA) and R01AI074345. The research on lymphatic filariasis on Mauke was supported by the Division of Intramural Research of the National Institute of Allergy and Infectious Diseases, National Institutes of Health. The original Haiti studies were supported by the United States Centers for Disease Control and Prevention (CDC), the National Institutes of Health, and the United Nations Development Program/ World Bank/ World Health Organization Special Program for Research and Training in Tropical Diseases (grant # 920528 and # 940441). KLH was supported by a CDC/APHL Emerging Infectious Disease Fellowship. The funders had no role in study design, data collection and analysis, decision to publish, or preparation of the manuscript.

**Measuring changes in transmission of neglected tropical diseases, malaria, and enteric pathogens from quantitative antibody levels**

**Table S1.**
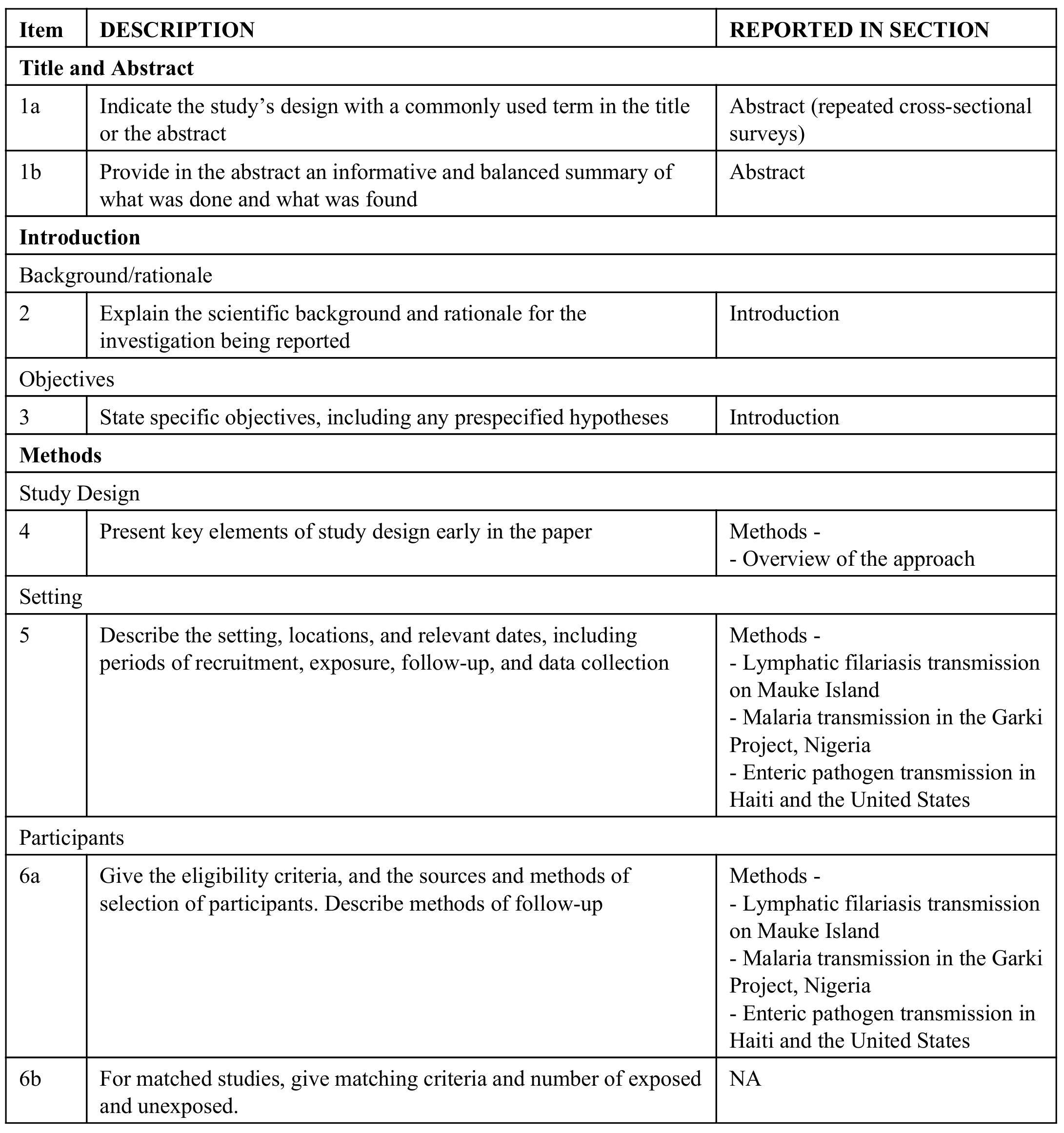
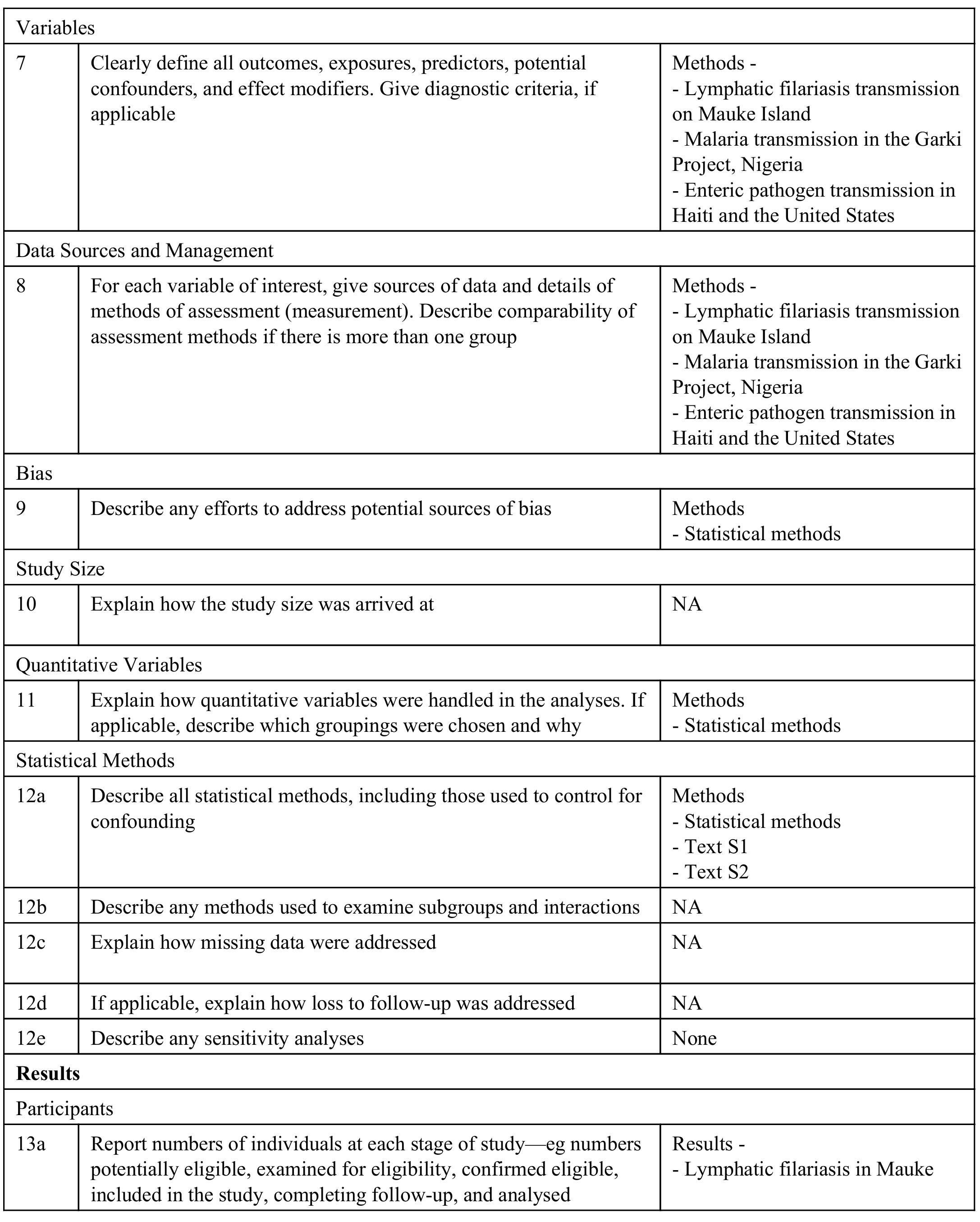
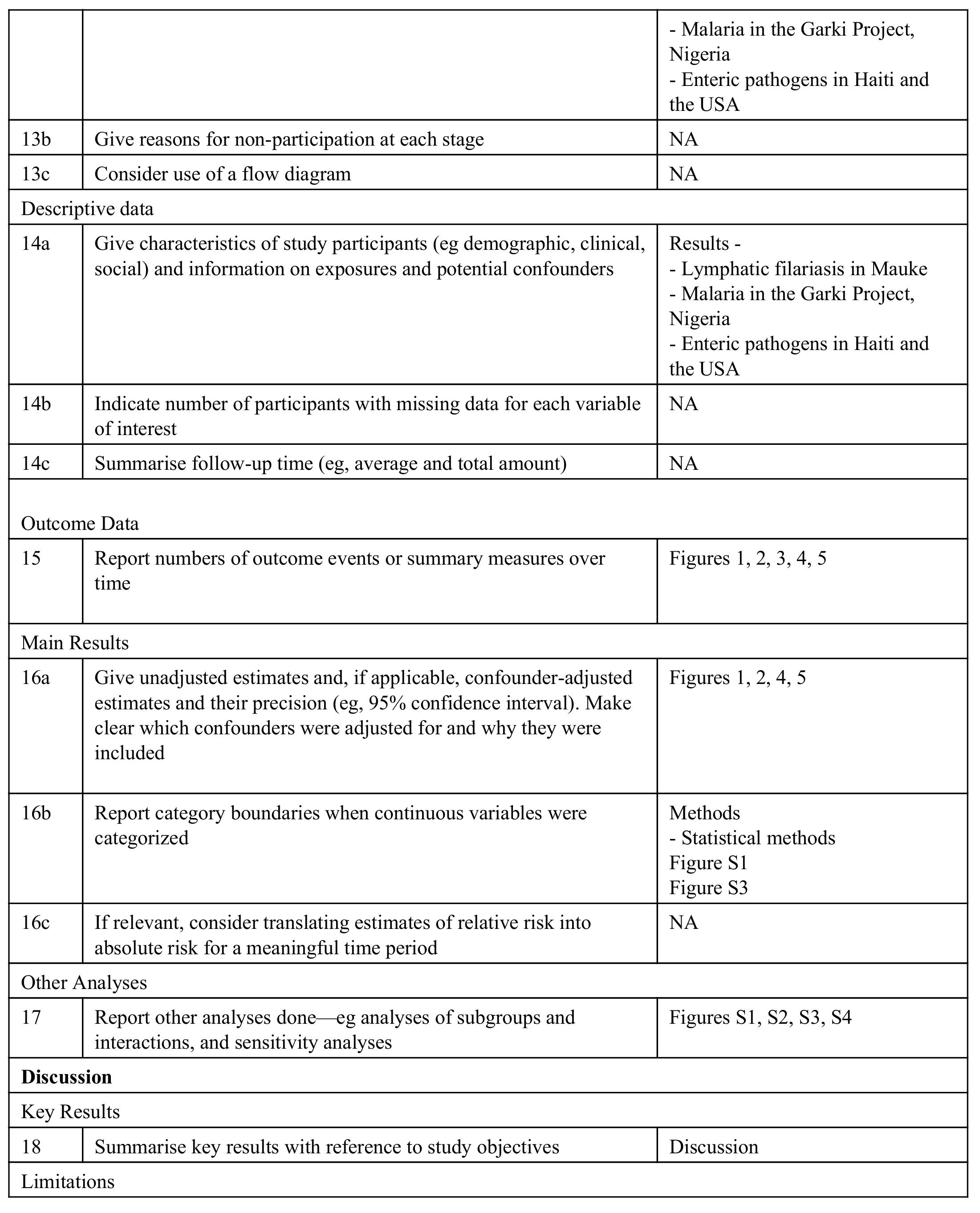
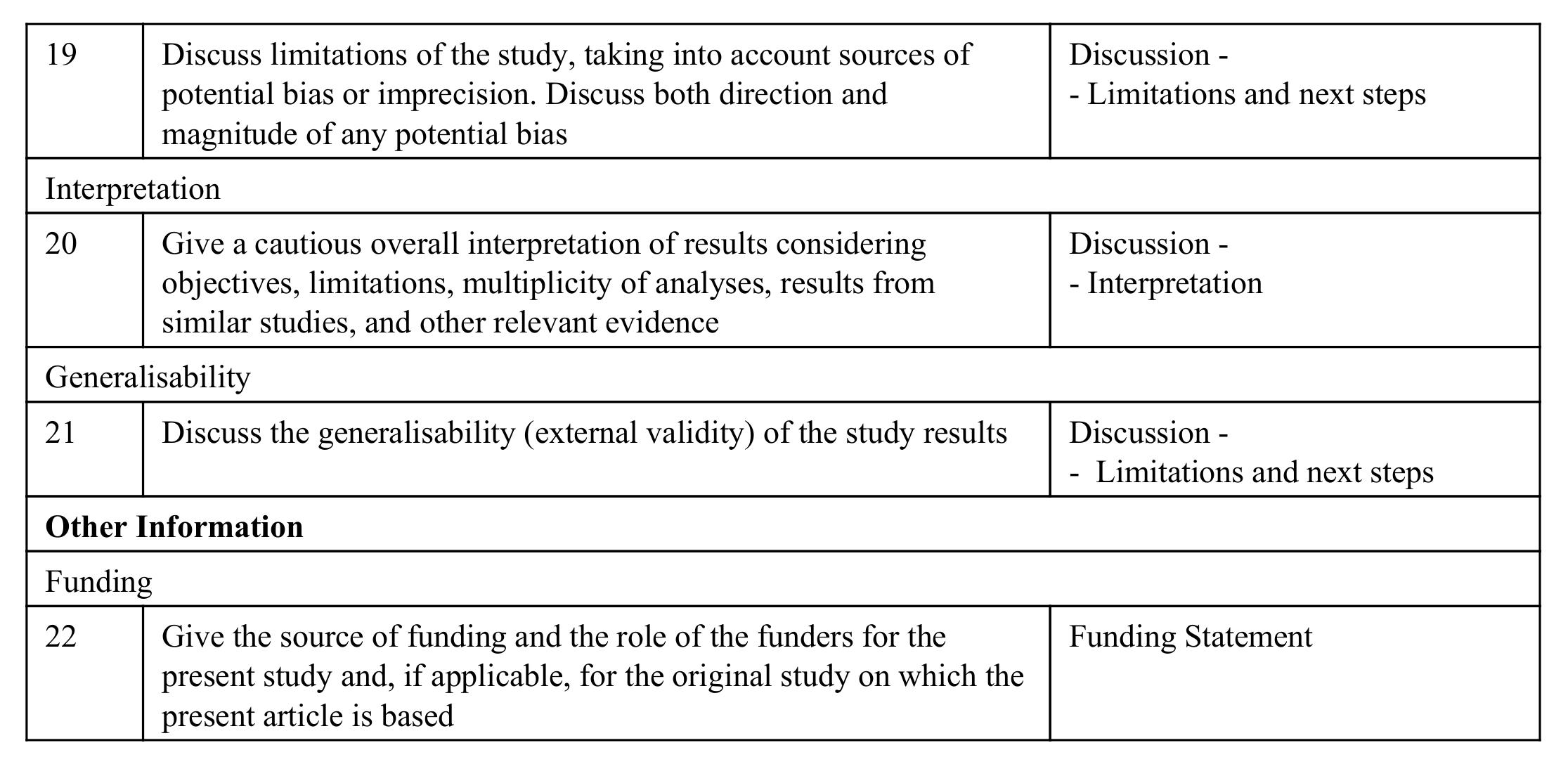
Strengthening the Reporting of Observational Studies in Epidemiology (STROBE) Checklist

**Measuring changes in transmission of neglected tropical diseases, malaria, and enteric pathogens from quantitative antibody levels**

**Text S1: Relationship between the age-adjusted mean antibody response and the area under the curve**

As in the main text Methods, the observed data on individuals include a quantitative antibody response (*Y*), age (*A*), a categorical exposure of interest (*X*), and a set of potentially confounding covariates (*W*). We observe *n* i.i.d. copies of *O* = (*Y*, *A*, *X*, *W*) with probability distribution *O* ~ *P*_0_. Age-antibody curves are the mean antibody response by age (*A* = *a*) and exposure (*X* = *x*), marginally averaged over *W*:

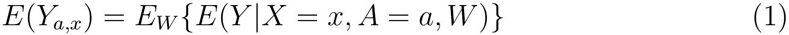

A smooth function of the age-antibody curve is the overall mean antibody level conditional on exposure group (*X* = *x*):

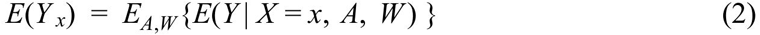

The following illustrates the relationship between this marginal mean and the area under the age-antibody curve:

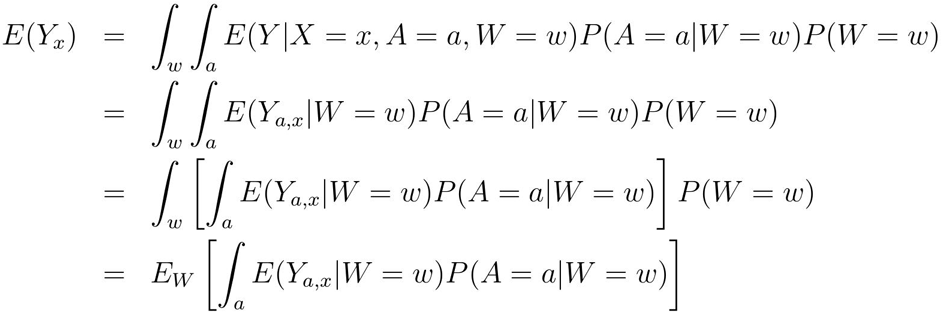

The term inside the brackets is area under the age-antibody curve (AUC), within strata defined by *W* = *w* and weighted by *P* (*A* = *a*|*W* = *w*). *E*(*Y*
_*x*_) is thus the marginal average across the straum-specific AUCs. In a special case where age is independent of other covariates, or investigators do not need to condition on potential confounders, *P* (*A* = *a*|*W*) = *P* (*A* = *a*) and the above expression further reduces to:

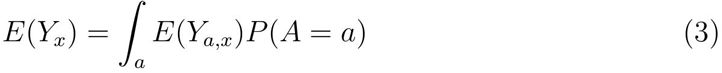

This quantity is the AUC of the age-antibody curve in exposure group *X* = *x*.

**Measuring changes in transmission of neglected tropical diseases, malaria, and enteric pathogens from quantitative antibody levels**

**Text S2: Technical details of estimating age-dependent antibody curves and changes in mean antibody levels**

Targeted maximum likelihood estimation (TMLE) is a double-robust, efficient estimation approach that targets the fit of the likelihood to the target parameter of interest [1, 2]. Below, we describe details of the estimation process and link the statistical estimation procedure directly to the observed data through a causal model [3]. TMLE is implemented in R using the tmle package (see [4] for a helpful overview of the package and methodology). A full set of replication files to implement the analyses described in this article, including a companion R package (tmleAb) complete with introductory vignette, are available through GitHub and the Open Science Framework: https://osf.io/8tqu4.

### Observed data and causal model

As in the main text Methods, a cross-sectional survey measures an individual’s quantitative antibody level (*Y*), age (*A*), and other characteristics (*W*). Many surveillance efforts are also interested in differences in antibody levels by one or more exposures (*X*), which may be confounded by *W*. We assume the observed data *O*= (*Y*, *A*, *W*, *X*) ~ *P*_0_ arise from a simple causal model:

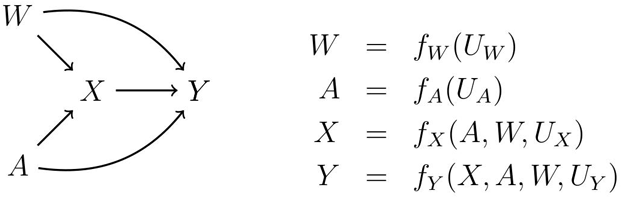

For simplicity, the graph on the left omits unmeasured characteristics (*U*) that, together with each variable’s parents, determine what value it takes (e.g., the error term for *X* is denoted *U*_*X*_). The equations encode assumptions about time-ordering between variables in the data (e.g., *X*, *A*, and *W* precede *Y*), but make no assumptions about the functional form of the relationship between them. The equations make no formal assumption about the relationship between errors that generate the data, but under the untestable assumption of no unmeasured common causes of *Y* and *X*, (independence of *U*_*Y*_ and *U*_*X*_), then conditional on *A* and *W* mean differences between groups defined by *X* have a causal interpretation.

### Fitting age-dependent antibody response curves

We are interested in a nonparametric model the mean antibody response as a function of age *A*, exposure *X* and potential additional covariates *W* (all defined above).

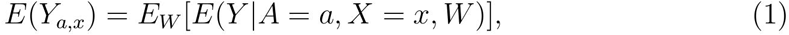

where the outside expectation marginally averages over the covariates *W*. A flexible approach to modeling the mean is to use an ensemble of models and algorithms to predict the mean over observed values of age and exposure. To accomplish this, we used the “super learner” algorithm, which is a stacked regression approach that combines individual predictions from each model or algorithm in its library into a single prediction that minimizes the cross-validated loss – in this case the mean squared error [5]. The algorithm is implemented in the SuperLearner package in R, and the companion R package for this paper (tmleAb) includes a convenient interface for flexibly estimating antibody curves.

In this analysis, we included in the ensemble: the simple mean, generalized linear models, antibody acquisition models with constant rates [6], locally weighted regression (lowess) [7], generalized additive models with natural splines [8], multivariate adaptive regression splines [9], and Random Forest [10]. We used default tuning parameters with two exceptions. First, for generalized additive models we selected the degrees of freedom smoothing parameter in natural splines for each fit using cross-validation [11]. Second, the default Random Forest implementation in R can overfit the data by growing trees that are too deep (i.e., have too small nodes), so for each fit we selected the minimum node size in Random Forest using cross-validation.

After fitting the ensemble, we generated curves by predicting the mean outcome for each level of age and exposure each analysis. Nonparametric estimates of an exposure-response curve with a continuous exposure (here: age) do not converge at a *n*^1/2^ rate [12] so pointwise confidence intervals will typically have poor coverage, even under a semi-parametric generalized additive model [13]. Below, we propose a summary function of the curve, which can be estimated consistently and efficiently under a nonparametric model.

### Target parameter of interest (estimand) to compare groups

Our parameter of interest of the observed data distribution *P*_0_ was the difference in the age- and covariate-adjusted population average antibody response in different groups. For example, in the Mauke study the exposed group (*X* = 1) included measurements after mass drug administration (MDA), and unexposed (*X* = 0) was pre-MDA:

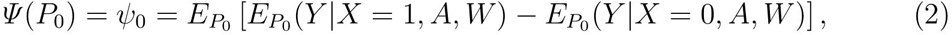

where the outer expectation is the mean averaged over the empirical distribution of *A* and *W*. *ψ* represents a mapping from a probability distribution *P*_0_ to a real number – namely, the difference in means. This target parameter depends on the probability distribution *P*_0_ of *O* through two quantities: the conditional mean, *E*_*P*0_ (*Y* |*X*, *A*, *W*), and the marginal distribution of (*A*, *W*). Due to the high dimensionality of the statistical model, standard maximum likelihood estimation, which maximizes the likelihood over all possible probability distributions, is not possible or results in overfitted estimators. Instead, targeted maximum likelihood (TMLE) is a two stage procedure that first uses ensemble machine learning to obtain an initial fit of these quantities and subsequently targets the fit so that it is optimal for the resulting plug-in estimator of *ψ*_0_.

### Targeted maximum likelihood estimation (TMLE)

For *n* observations from the probability distribution *O* ~ *P*_0_, we define a plug-in estimator of *ψ*_0_ with the form:

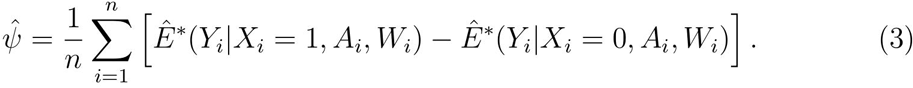

TMLE estimates 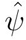 using a two-stage process. The first stage makes an initial estimate of *Ê*^*^ (*Y*|*X*, *A*, *W*), denoted *Ê*^0^ (*Y*|*X*, *A*, *W*). This could be done using a highly parametric approach, such as linear regression, but the relationship between antibody response, exposure group, age, and other characteristics could be very complex. As with the estimation of age-dependent antibody curves, we used the “super learner” algorithm to flexibly estimate *Ê*^0^(*Y*|*X*, *A*, *W*), which are predicted antibody levels conditional on *X*, *A*, and *W*.

The second stage of TMLE updates the the initial fit, *Ê*^0^(*Y*| *X*, *A*, *W*) is consistent then the TMLE targeting step. If the initial estimate of E estimate is consistent, but if it is biased then the updating step helps remove residual bias in the estimation of *ψ*_0_. It involves first estimating a nuisance parameter: the probability of treatment given observed characteristics, sometimes called the propensity score [14]. In randomized trials *P* (*X* = *x*|*A*, *W*) is known, but in observational studies *P* (*X* = *x*|*A*, *W*) is typically not known and must be estimated. We estimated 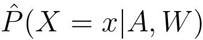 using the super learner algorithm with the same library of learners described above, but with *X* as a binary outcome variable. The analysis assumes that *P*_0_(*X* = 1|*A* =*a*, *W* = *w*) > 0 and *P*_0_(*X* = 0|*A* = *a*, *W* = *w*) > 0 are positive. Without this assumption, the conditional expectations of Y in *Ψ* (*P*_0_) are not well defined. In practice, this second assumption means that exposure groups need to have good overlap in age and any other covariates included in the analysis.

TMLE then uses 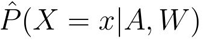 to construct an individual-level covariate, which is chosen specifically for our parameter of interest (equation 2) to solve the efficient influence curve and thus minimize the bias in the estimate of *ψ*_0_[2]:

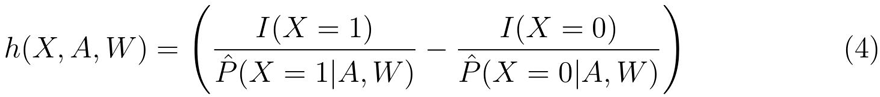

The targeted update is achieved using a univariate generalized linear model that includes *Ê*^0^(*Y*|*X*, *A*, *W*) as the offset with *h*(*X*, *A*, *W*) as a single covariate. For continuous outcomes such as *Y*, it has been shown that a conducting the update on the linear scale works in many cases (i.e., a generalized linear model with an identity link), but if *h* (*X*, *A*, *W*) can take on large values then there are important robustness advantages to conducting the update on the logistic scale using a re-scaled version of the outcome bound between (0,1) [15]. If *Y* is bounded by (*a*, *b*) then a re-scaled version *Y*^*^= (*Y* − *a*)/(*b* − *a*) is bounded by (0,1). The logistic model is then fit using maximum likelihood:

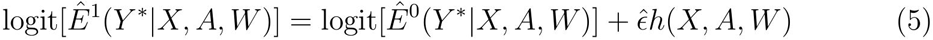

Predicted antibody levels from this update, transformed back to their original scale, are then used in the plug-in estimator: *Ê*^1^(*Y*|*X*, *A*, *W*) = *Ê*^*\**^ (*Y*|*X*, *A*, *W*) in equation 3. TMLE is a regular, asymptotically linear estimator. For this reason, standard errors, confidence intervals, and *P*-values are estimated using the influence curve [1, 2], and readily accommodate repeated measures data if that is a feature of the design [4].

